# Post-meiotic mechanism of facultative parthenogenesis in gonochoristic whiptail lizard species

**DOI:** 10.1101/2023.09.21.558237

**Authors:** David V. Ho, Duncan Tormey, Aaron Odell, Aracely A. Newton, Robert R. Schnittker, Diana P. Baumann, William B. Neaves, Morgan R. Schroeder, Rutendo F. Sigauke, Anthony J. Barley, Peter Baumann

## Abstract

Facultative parthenogenesis (FP) has historically been regarded as rare in vertebrates, but in recent years incidences have been reported in a growing list of fish, reptile, and bird species. Despite the increasing interest in the phenomenon, the underlying mechanism and evolutionary implications have remained unclear. A common finding across many incidences of FP is a high degree of homozygosity at microsatellite loci. This has led to the proposal that first or second polar body fusion following the meiotic divisions restores diploidy and thereby mimics fertilization. Here we show that FP occurring in the gonochoristic *Aspidoscelis* species *A. marmoratus* and *A. arizonae* results in genome-wide homozygosity, an observation inconsistent with polar body fusion as the underlying mechanism of restoration. Instead, a high-quality reference genome for *A. marmoratus* and analysis of whole-genome sequencing from multiple FP and control animals reveals that a post-meiotic mechanism gives rise to homozygous animals from haploid, unfertilized oocytes. Contrary to the widely held belief that females need to be isolated from males to undergo FP, females housed with conspecific and heterospecific males produced unfertilized eggs that underwent spontaneous development. In addition, a mixture of offspring arising from fertilized eggs and parthenogenetic development was observed to arise from a single clutch. Strikingly, our data support a mechanism for facultative parthenogenesis that removes all heterozygosity in a single generation. Complete homozygosity exposes the genetic load and explains the high rate of congenital malformations and embryonic mortality associated with FP in many species. Conversely, FP constitutes strong purifying selection as non-functional alleles of all essential genes are purged in a single generation.

## Introduction

Incidences of facultative parthenogenesis (FP) have been reported to occur in diverse vertebrate clades including bony fish (1), sharks (2–5), snakes (6–12), lizards (13–15), crocodilians (16), and birds (17–21). The phenomenon was originally mistaken for long-term sperm storage occurring in zoo environments where females were housed without current or recent access to conspecific males. The most parsimonious explanation was therefore that the animal had previously been in contact with a male and that stored sperm was responsible for delayed fertilization (8, 22, 23). However, more recent studies involving microsatellite (MS) and/or amplified fragment length polymorphism (AFLP) analyses revealed no paternal contributions, as all alleles detected in the offspring were also found in the maternal ancestors (7, 8, 24, 25). Females with no access to males producing solely male (ZW systems) or female (XY systems) offspring that only harbor maternal genetic markers are now considered hallmarks of facultative parthenogenesis. Originally thought to only occur in captivity, more recent reports indicate that FP occurs in natural populations as well (10, 26). Serious concerns have been raised by conservation biologists, as species with dwindling population densities, including the critically endangered species Komodo dragon (14), small tooth sawfish (26), and California condor (25), are overrepresented among reports of FP. While overrepresentation could be a consequence of an increased likelihood of detection in species that are the subject of intense research and conservation efforts, the observations are also consistent with the hypothesis that FP is an adaptive trait aiding in the colonization of new areas and mitigating the effects of population bottlenecks when mate encounters become less frequent (26). At the same time, the association of FP with increased homozygosity constitutes a considerable concern for conservation biology, as an increase in FP within dwindling populations further accelerates the loss of genetic diversity and compromises efforts to maintain the existing gene pool in selective breeding programs (7, 27). In another context, FP is being studied as a desirable outcome in the commercial production of poultry. However, examination of tens of thousands of unfertilized eggs from several different avian species and strains has not resulted in economically viable hatching rates thus far. One of the highest hatch rates for unfertilized eggs is seen in the Beltsville small white turkey with a rate of 0.88% (21, 28). A better understanding of the triggers and molecular mechanisms underlying FP is therefore urgently needed to aid in conservation efforts including captive breeding programs and to possibly harness FP in an agricultural context (25).

The high level of homozygosity observed in animals produced by FP has been interpreted as evidence for polar body fusion following meiosis II, also known as automixis, leading to the restoration of diploidy in unfertilized eggs (Fig. 1*A*) (2, 16, 27, 29). If automixis involves the fusion of one of the meiotic products from the first polar body (central automixis), homozygosity will be concentrated near the chromosome ends and heterozygosity will be preferentially retained near the centromeres as premeiotic recombination strongly favors homologs over sister chromatids and homologs segregate during the first meiotic division. In contrast, a second polar body fusion (terminal automixis) would reunite sister chromatids, for which heterozygosity is preferentially seen near the chromosome termini (Fig. 1*B*). In some cases, heterozygous and homozygous loci appeared to be inherited in FP offspring (7, 15), but information as to the genomic location of these loci has been lacking. Most FP case studies examined a small number of genetic markers, and homozygosity was suspected or confirmed for every maternally contributed allele, an observation that is also consistent with a post-meiotic mechanism of genome duplication that would produce genome-wide homozygosity (24). In aggregate, sparsity of available data has prohibited conclusive determination of the mechanism of FP in any vertebrate system thus far, despite the realization of an increasingly widespread occurrence.

**Figure 1.**
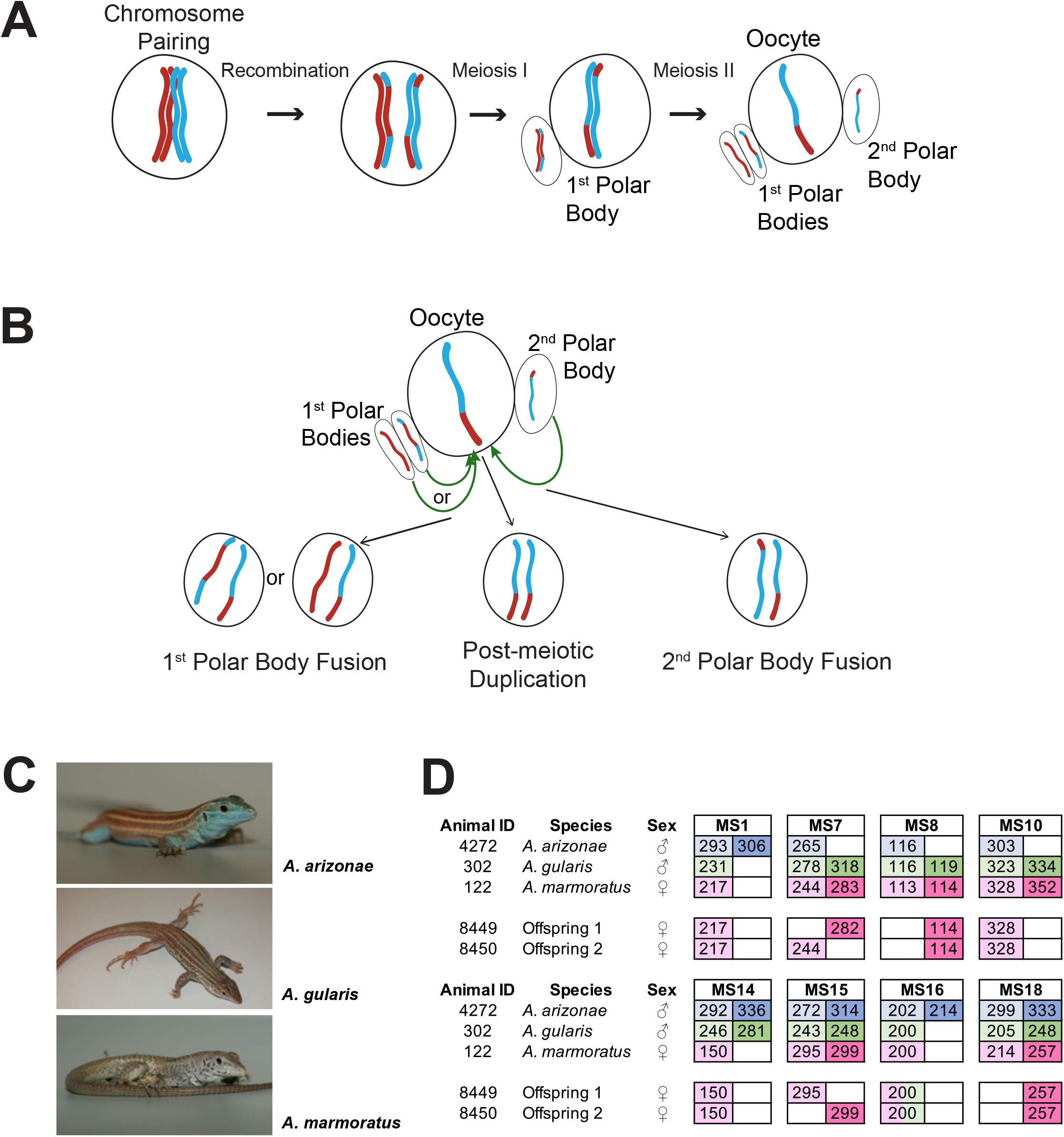
Overview. **(A)** Schematic of canonical meiosis. Only one pair of homologous chromosomes is shown using red and blue to distinguish homologs. **(B)** Schematic of mechanisms by which a diploid oocyte may be produced in the context of facultative parthenogenesis. First bolar body fusion, second polar body fusion or post-meiotic duplication of chromosomes in the haploid gamete. **(C)** Photographs of *Aspidoscelis arizonae* (top), *A. gularis* (middle), and *A. marmoratus* (bottom). **(D)** Microsatellite analysis for the three co-housed animals and two offspring (ID 8449 and 8450) produced in this enclosure. Alleles are color coded as follows: *A. arizonae* male (blue), *A. gularis* male (green) and *A. marmoratus* female (red). Differences in shading highlight the two alleles at heterozygous loci. Both offspring are homozygous at all loci with most alleles matching maternal alleles. For MS16 offspring alleles are shown in red and green to indicate the theoretical possibility of this allele being either inherited from the mother or the *A. gularis* male.

While facultative parthenogenesis occurs in a wide range of vertebrate species, true obligate parthenogenesis is limited to a few taxa of squamate reptiles including the North American whiptail lizards of the genus *Aspidoscelis* (30). Historic hybridization events between distinct gonochoristic species gave rise to hybrid individuals with the ability to reproduce clonally as all female lineages (31, 32). In contrast to the increased homozygosity associated with FP, obligate parthenogenetic species are characterized by the long-term preservation of the high degree of heterozygosity that had its origin in the lineage-founding cross species hybridization events.

Our laboratory has a longstanding interest in the mechanism of obligate parthenogenesis in whiptail lizards (33, 34). In this context we are housing and propagating individuals of several obligate parthenogenetic species as well as gonochoristic sister species. A systematic MS analysis for all individuals of gonochoristic species produced in our colony revealed over 20 incidences of FP in the marbled whiptail lizard *A. marmoratus* and the Arizona striped whiptail *A. arizonae*\*. Whole-genome sequencing of five individuals revealed genome-wide homozygosity for all FP animals and identified post-meiotic genome duplication, rather than polar body fusion, as the mechanism of FP in gonochoristic whiptail lizards. Examination of peripheral blood revealed mixoploidy in erythrocytes from FP animals, suggesting that genome duplication occurs during embryonic development rather than preceding it. Furthermore, we observed reproduction by FP in females in the presence of heterospecific or conspecific males. Arguing against a developmental switch separating sexual reproduction from FP, a single clutch can be comprised of fertilized and unfertilized eggs, with the unfertilized eggs undergoing complete FP development. To address the question whether FP is limited to animals in captivity, we examined reduced-representation sequencing data of 321 whiptail lizards from 15 gonochoristic species sampled in nature. Heterozygosity analysis identified several candidates of FP origin based on the extent of homozygosity. In aggregate, these findings reject the idea of a reproductive switch that activates parthenogenesis only in the absence of mating opportunities and suggests that a baseline incidence of FP may coexist alongside sexual reproduction in some species.

## Results

### Identification of FP in *A. marmoratus*

In the context of studying interspecific hybridization among gonochoristic species of whiptail lizards, three female *A. marmoratus* (ID 122, 4238, 4239) were housed with a male *A. arizonae* (ID 4272) and a male *A. gularis* (ID 302) for close to three years. During this period, seven hybrid offspring between *A. marmoratus* 122 and *A. arizonae* 4238 were produced and confirmed by MS analysis. These animals will be described in more detail in due course. Surprisingly, two female hatchlings emerged that resembled *A. marmoratus* rather than the expected products of hybridization with either *A. arizonae* or *A. gularis*. Genotyping revealed only a single allele for each of eight MS markers in the two offspring (ID 8449 and 8450, Fig. 1*D*) and identified *A. marmoratus* (ID 122) as the mother (Fig. S1). The mother and the male *A. arizonae* were each heterozygous at five of the eight markers and the *A. gularis* male at six. Further supporting a uniparental origin of 8449 and 8450, all alleles found in the offspring were also present in the mother (Fig. 1*D*). For seven of the eight markers, neither male shared the allele found in the hatchling lizards. For the remaining marker MS16, the *A. marmoratus* mother and the *A. gularis* male were homozygous for the same allele found in the two offspring. It is further important to note that for two of the markers (MS7 and MS15), the two offspring inherited different alleles from the mother, indicating that they are not genetically identical to each other, but have randomly inherited one of the maternal alleles at each locus.

Two additional eggs (ID 8394 and 9070) were recovered from the same enclosure and found to contain developing embryos. MS analysis also revealed *A. marmoratus* 122 as the mother and complete homozygosity at all loci supported FP origin. The inheritance of only maternal alleles strongly suggested FP rather than interspecific hybridization or long-term sperm storage.

FP animals of *A. marmoratus* presented a unique opportunity to examine the underlying molecular mechanism. The observed homozygosity at all MS loci further promised to aid in the generation of a high-quality genome assembly, as homozygosity circumvents the challenge of collapsing haplotypes into a consensus sequence (37). In order to increase homozygosity, inbreeding for 15 to 20 generations is common practice prior to whole-genome sequencing and genome assembly (38, 39). However, generation times of more than one year make this a costly and time-consuming strategy for *A. marmoratus* and many other vertebrates. As facultative parthenogenesis has been reported to consistently generate a high degree of homozygosity in a single event, such animals provide a good starting material for de novo genome assemblies.

### Genome sequencing and *de novo* assembly

The *A. marmoratus* genome is distributed over 23 chromosomes as previously demonstrated by metaphase spread analysis (40). We used flow cytometry to compare nuclear DNA content of *A. marmoratus* erythrocytes from whole blood with cells from three species with well-characterized genome sizes. The nuclear DNA content of *A. marmoratus* was close to that of *Danio rerio* and we calculated a haploid genome size for *A. marmoratus* of 1.67 Gb (Fig. S2*A*).

Genomic DNA of FP animal 8450 was used to generate short insert paired-end, mate-pair (5 Kb, 8 Kb, 2-15 Kb, 40 Kb), and Chicago (41) libraries for Illumina short-read sequencing. The paired-end and mate-pair reads were first assembled with Meraculous (42) yielding an N50 of 1.6 Mb. The subsequent addition of Chicago reads and scaffolding with the HiRise pipeline by Dovetail Genomics produced an assembly of 1,639,530,780 bp distributed over 3,826 scaffolds (Table S1) and raised the scaffold N50 to 32.22 Mb (Fig S2*B*). With a BUSCO completeness score of 96% the *A. marmoratus* genome assembly is comparable to other recently released reptilian genome assemblies (Fig. S2*C*). Over 98% of the assembled sequences are contained within 90 scaffolds of more than 1 Mb in length, making this assembly highly contiguous.

Phylogenetic analysis of shared BUSCO genes with several other reference genomes (*Xenopus*, zebrafish, medaka, platyfish, tegu, green anole, chicken, mouse, rat, dog, cow, human) confirmed that *A. marmoratus* is most closely related to the tegu *Salvator merianae*, another representative of the family Teiidae (Fig. S3). As transposable elements are a driving force in genome evolution, we examined the repeat content for the *A. marmoratus* genome. All classes of repeat elements combined amounted to 40.27% of the *A. marmoratus* assembly, only slightly below that found in other lizards *S. merianae* and *Anolis carolinensis* (Fig. S4*A*). Strikingly, unclassified repeats make up the largest class of repeat elements in the *A. marmoratus* genome, an observation that parallels findings in *S. merianae*. However, a comparison between the unclassified repeats found in *A. marmoratus*, *S. merianae,* and *A. carolinensis* revealed few similarities with only around 10% of the unclassified repeats shared between *A. marmoratus* and *S. merianae*, and no significant overlap between these two lizards and *A. carolinensis* (Fig. S4*B*). While further characterization of the unclassified repeat elements is needed, it is apparent that an impressive expansion of novel repeat element classes has occurred within this clade.

To annotate the *A. marmoratus* genome, we assembled a total of 119,728 transcripts from RNA-seq data generated from blood and embryo using Trinity (Table S2). These transcripts were subsequently used in the MAKER2 gene annotation pipeline, yielding 25,856 protein-coding genes and 44,461 protein isoforms (Table S3). To assign putative function, we used BLASTp to query the UniProtKB/Swiss-Prot database (43) and found significant hits for 76% of the putative protein-coding genes. Our assembly and annotation pipelines yielded 40 *HOX* genes and 2 *EVX* genes in four gene clusters (Fig. S5). The *HOX* gene clusters are highly conserved among tetrapods and their complete presence and shared order within each cluster serves as a measure of assembly quality (44).

### Assessment of heterozygosity

The presence of only one allele for each of the examined MS markers already suggested widespread homozygosity in *A. marmoratus* produced by FP, consistent with similar observations in other vertebrate species. The highly contiguous genome assembly now afforded us the opportunity to probe the mechanism of FP by searching for regions of heterozygosity and mapping their relative genomic locations. Towards this aim, we performed whole-genome sequencing for an additional nine animals: four of them produced by FP (ID 12512, 12513, 6993, 9177), two mothers (ID 122, 9721), and three unrelated control animals (ID 003, 001, S30700; Table S4). Each mother and the controls were heterozygous at several MS markers confirming their origin through sexual reproduction. Following alignment to the reference genome (Table S5), we defined heterozygous sites, as those having equal support for two alleles.

For all ten individuals, the average sequencing coverage ranged between 15.91 and 20.08 (Fig. S6). For FP animals, the number of heterozygous sites in a 10 kb sliding window approaches zero for all sites with mean coverage (Fig. 2*A*). For positions with coverage greater than the average, an increase in apparent heterozygosity was observed, due to the collapse of repetitive sequences during the assembly process. Based on this observation, we limited further analysis to positions in the genome where the coverage is equal to the mean sequencing depth. This generated between 30,769 and 53,416 heterozygous sites for which two alleles were equally supported in the sexually produced mothers and control animals (Fig. S7*A-E*). Far fewer heterozygous sites (between 649 and 928) were observed in the FP animals (Fig. S7*F-J*). Plotting the heterozygous sites according to their position in the reference genome illustrates not only their sparsity in the genomes of FP animals, but also reveals their random distribution (Fig. 2*B*). If FP genomes were the product of automixis, regions of homozygosity would be interspersed with regions of heterozygosity. The extent of heterozygosity within the latter would be the same as that observed in the respective mother. The few apparently heterozygous sites identified in FP animals are therefore not supporting either form of automixis but are most likely the result of a combination of replication, sequencing, and over-assembly of repetitive regions.

**Figure 2.**
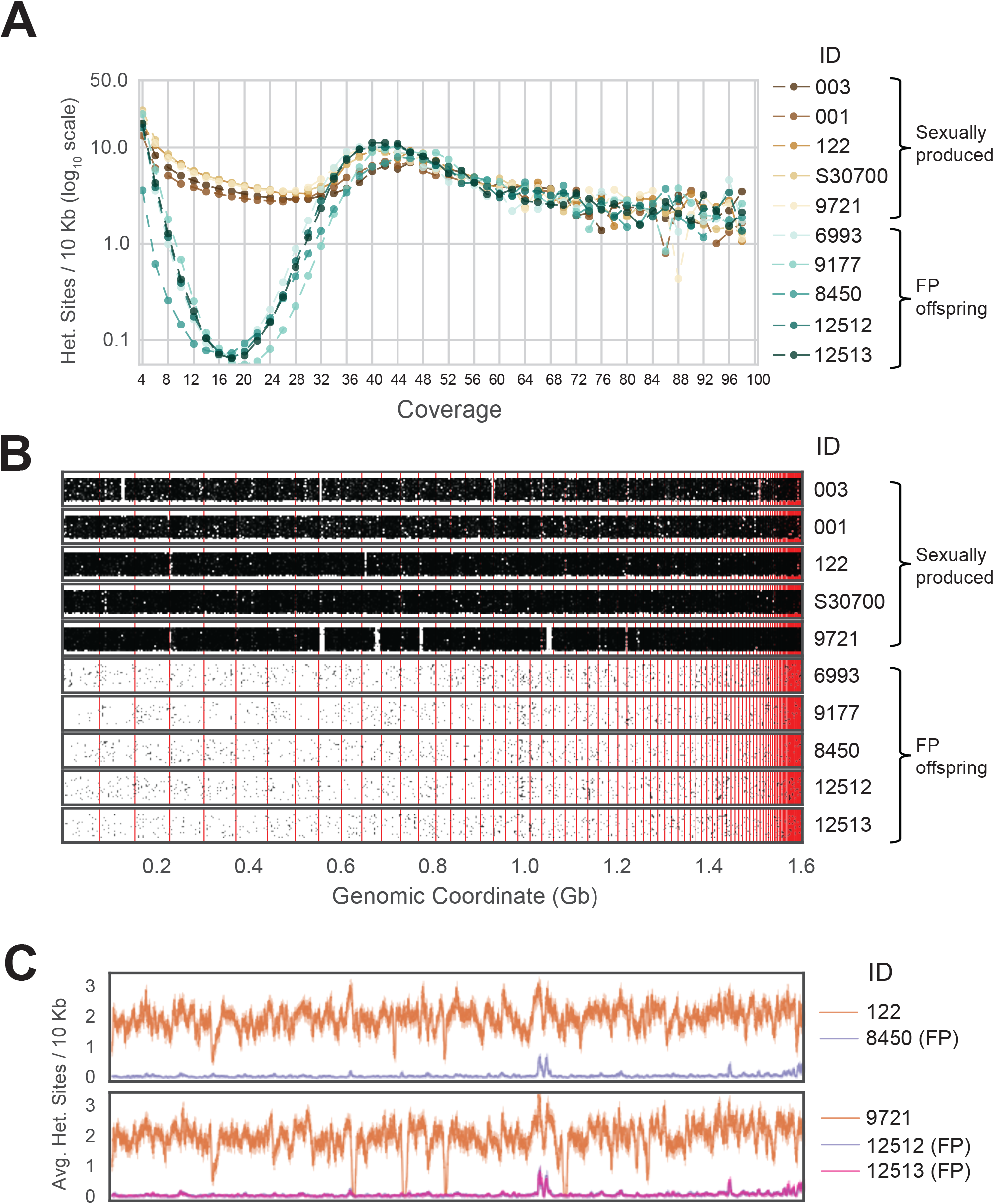
Genome-wide homozygosity in animals produced by facultative parthenogenesis. **(A)** Effect of coverage on the apparent rate of heterozygosity based on evenly split read-counts supporting two alleles. Analysis of whole-genome sequencing data for five sexually produced animals (ID 003, 001, 122, S30700, 9721) and five individuals produced by facultative parthenogenetic (FP) animals (ID 6993, 9177, 8450, 12512, 12513) were aligned to the reference genome. In only FP animals, the number of heterozygous sites approaches zero for sites with mean coverage (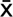 = 18.37). **(B)** Scaffolds are ordered from largest to smallest on the x-axis. Red lines indicate borders between ordered scaffolds. Each black dot represents a heterozygous position in the genome defined having a sequencing coverage equal to the average and equal support for only two alleles. The y-axis position is random value between bounds of area shown. **(C)** Average heterozygous sites, as defined in (B), per 10 Kb step for mother (orange) and FP daughter combinations (purple and pink).

When examining each mother-daughter group, the average number of heterozygous sites per 10 Kb window was greater in the sexually produced mothers across the entire assembly (Fig. 2*C*). For most of the assembly, the average number of heterozygous sites remained close to zero for FP animals. The most notable exception was a region around genomic coordinate 1.0 Gb, but even there the extent of apparent heterozygosity remained below 50% of what is observed in this region for each of the mothers. Examination of the scaffold in question (Scaffold 45) revealed 167 genes annotated as homologous to *vomeronasal 2 receptor 26* (*Vmn2r26;* Fig. S8, Fig. S9). Members of this subfamily of receptors are found on the microvilli of the vomeronasal organ, where they are responsible for pheromone detection and play a significant role in social and environmental responses (45–47). Given that this genomic region harbors a large cluster of highly similar genes, the most parsimonious explanation for the elevated level of apparent heterozygosity is over-assembly. This conclusion is further supported by the increase in apparent heterozygosity in this region for the mothers and control animals. In aggregate, our analysis strongly supports genome-wide homozygosity for FP animals, inconsistent with either central or terminal automixis. Instead, the results favor a post-meiotic mechanism that restores diploidy by replicating the haploid genome residing in the oocyte following completion of the two meiotic divisions and thereby establishing genome-wide homozygosity in the offspring.

### Cryptic FP in a mixed clutch

Following the identification of several *A. marmoratus* generated by FP, we genotyped all individuals from two other gonochoristic species housed in our laboratory. While no cases of FP were identified among 80 *A. gularis* produced in captivity, we identified eight incidences of FP among 832 *A. arizonae* hatched between October 2007 and July 2018. During the same period, we recorded 15 incidences of FP among 286 *A. marmoratus* (Table S6). Notably, in all cases, eggs undergoing FP development had been laid in enclosures where females were housed with conspecific males or males of a sister species known to mate with the heterospecific females. Isolation from mating partners was thus not a significant factor in triggering FP. In one enclosure, three *A. arizonae* females (ID 12850, 12851, 12852) were housed with a conspecific male (ID 12849; Fig. 3*A*). MS analysis of four hatchlings that originated from a single clutch laid in this enclosure identified animal 12852 as the mother of all four animals. Unexpectedly, two of her offspring were homozygous at all eight loci examined, whereas the two others were heterozygous at all loci, identifying animal 12849 as their father (Fig. 3*B*). Therefore, both fertilized and unfertilized eggs developed alongside each other within the same clutch.

**Figure 3.**
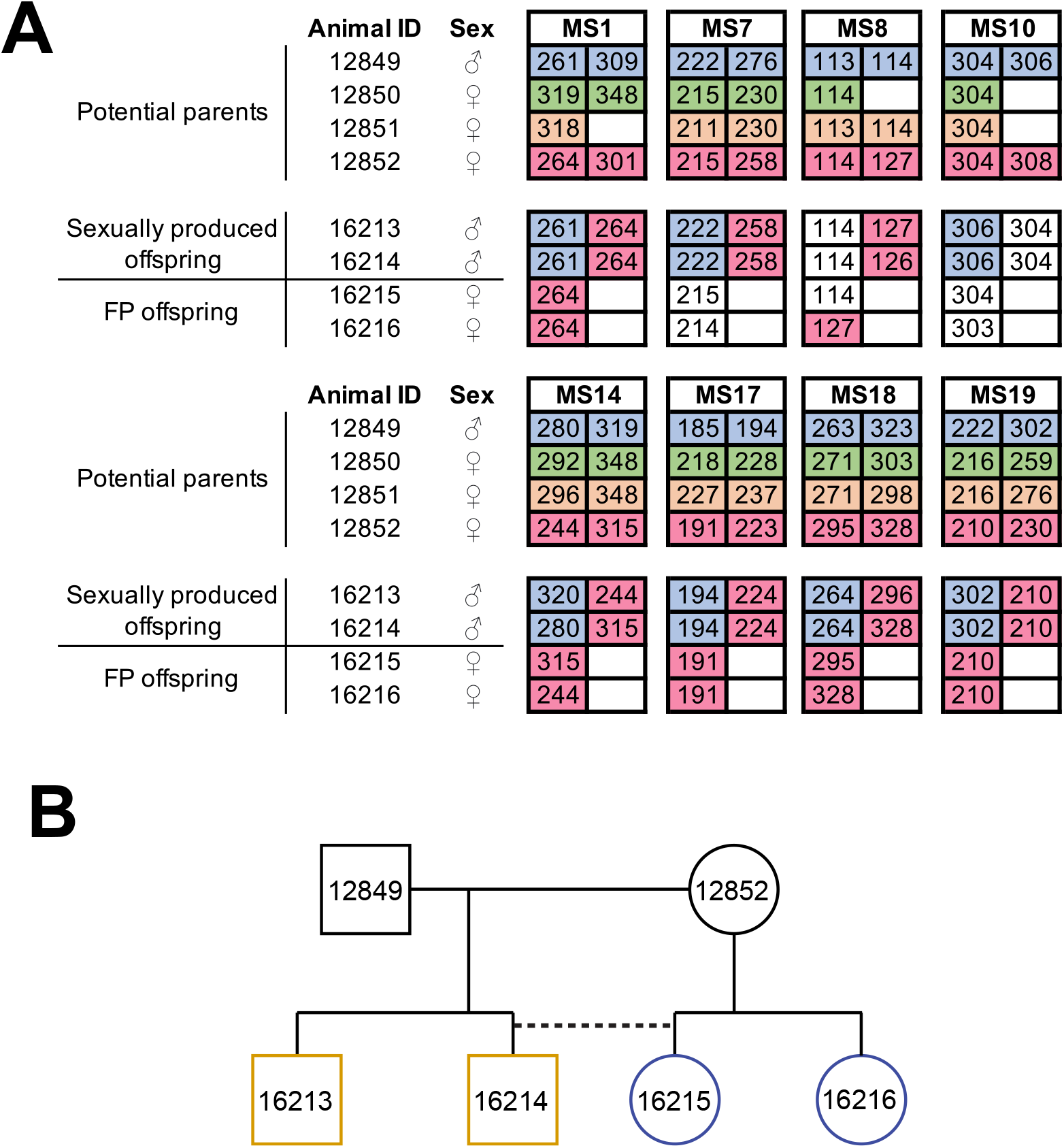
Facultative parthenogenesis is also found in *Aspidoscelis arizonae*. **(A)** Microsatellite analysis for the four co-housed animals and the four offspring (ID 16213, 16214, 16215, 16216) produced in this enclosure. Alleles are color coded for each potential parent: 12849 male (blue), 12850 female (green), 12851 female (orange), 12852 female (red). Offspring 16213 and 16214 are heterozygous at all loci, with most loci having one allele matching 12849 and one allele matching 12852. Offspring 16215 and 16216 are homozygous at all loci, with most alleles matching only the 12852 female. **(B)** Pedigree showing the relationship between the four offspring. The single clutch of four contains both sexually (yellow) and facultative parthenogenetically (blue) produced offspring.

### Mixoploid erythrocytes and developmental defects

Microscopic examination of blood from newly hatched FP lizards revealed a striking bimodality in the sizes of erythrocyte nuclei (Fig. 4). Nuclear size correlates well with DNA content measurements (48), suggesting the presence of mixoploidy in these animals. Whereas most cells closely resembled those observed in the blood from sexually produced animals, approximately 10% of red blood cells from FP animals harbored smaller nuclei, consistent with half the amount of DNA (Fig. 4*B*). In addition, 1.27% contained two small nuclei, indicating that the final cytokinesis during erythrocyte differentiation had failed for some haploid progenitor cells. These observations raise the possibility that embryonic development initiated with consecutive divisions of a haploid, unfertilized oocyte. At a later stage in development, diploid cells most likely arise via failed cytokinesis. From that point forward, both haploid and diploid cells coexist, and the embryo develops in a mixoploid state. Indicative of a more widespread phenomenon, mixoploidy was also observed in *A. arizonae* (Fig. S10).

**Figure 4.**
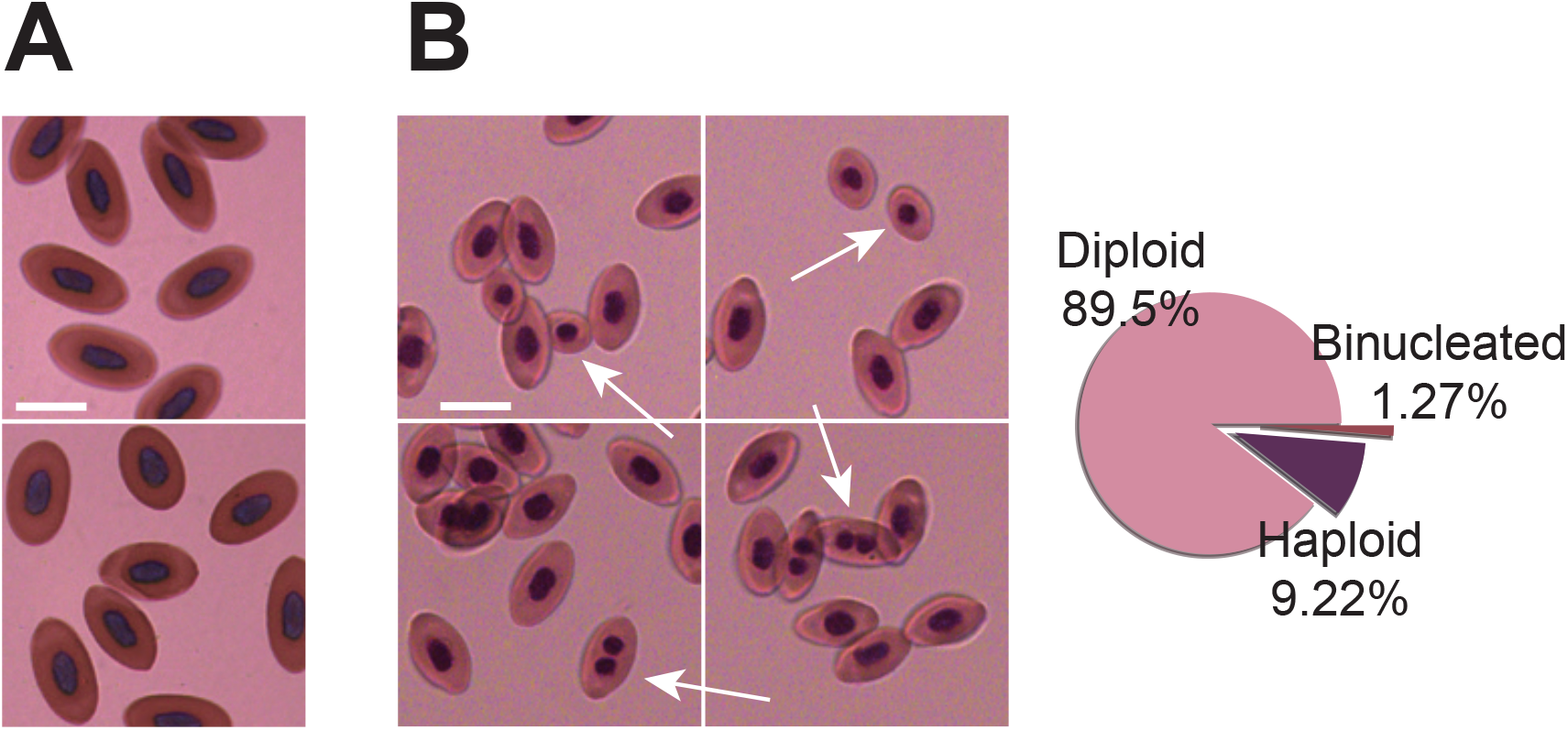
Animals produced by facultative parthenogenesis are characterized by mixoploidy. **(A)** Giemsa staining of erythrocytes from a sexually produced *Aspidoscelis marmoratus*. All cells are diploid (n = 601). Scale bar is 10 µm. **(B)** Giemsa staining of erythrocytes from a newly hatched facultative parthenogenetic *A. marmoratus*. Diploid (n = 844), smaller haploid (n = 87), and binucleated (n = 12) cells are evident. Scale bar is 10 µm.

Genome-wide loss of heterozygosity exposes functionally compromised alleles that were previously covered by intact alleles on the homologous chromosomes. Depending on the extent of this genetic load, one would expect a substantial fraction of oocytes to not develop at all or for defects to manifest at various stages of embryonic and post-embryonic development. Indeed, of the 23 incidences of FP examined here, only 16 hatched, while the remaining lizards died in ovum (Table S6). For these seven unhatched eggs, we isolated developed embryos shortly after the expected hatch date and confirmed FP origin by MS analysis. The clutches that produced the 23 confirmed cases of FP contained an additional 24 eggs. For these, development did not initiate or terminated at an early stage of development precluding MS analysis. Despite the uncertainty regarding how many of these eggs underwent partial FP development, the incidence of FP may be even higher than reported here.

The observation of various malformations in several of the FP embryos and hatchlings further supported that genome-wide homozygosity unmasks deleterious alleles. Notable developmental defects included craniofacial abnormalities such as misaligned jaws and agenesis of eyes, missing limbs, and failed abdominal closure (Fig. S11). In contrast, six FP animals hatched with no discernable developmental defects (Fig. S11*A-B*). While we have not recorded instances of FP animals producing offspring via FP, as described for the whitespotted bambooshark (49), FP *A. marmoratus* 8450 did produce two eggs while housed in isolation that did not hatch. Analysis of the ovaries of FP animal 8450 as well as germinal vesicles of its FP sister 8449 revealed no differences in structure and anatomy compared to fertile sexually reproducing animals (Fig. S12).

### Incidence of FP in whiptail lizards

As FP has been associated with captivity in most species where it has been reported, we examined restriction site-associated DNA sequencing (RAD-seq) data for wild animals across 15 gonochoristic species (Dataset S1; Fig. S13). Because RAD-seq is a form of reduced-representation sequencing (50), we limited further analysis to the 321 individuals that had an average sequencing coverage of at least 20. Computational data analysis revealed five animals that had very low levels of heterozygosity at positions of average coverage (< 0.05% heterozygous positions). In contrast, the average level of heterozygosity was 0.261% across the dataset. Of the five low heterozygosity animals, *A. deppii* (ID LDOR30) showed the most striking level of homozygosity affecting all sites but one (Rosner’s Test for Outliers, log-transformation, *R* = 5.127, λ = 3.928, *p* < 0.001). This is consistent with the pattern of homozygosity observed with the whole-genome sequencing from FP animals produced in the laboratory. Further field work and analysis will be required to assess the level of FP in natural populations of gonochoristic *Aspidoscelis* species, but analysis of the available dataset puts the rate between 0.3% and 1.5%. This is a similar range to the 1% and 5% observed in captive *A. arizonae* and *A. marmoratus*, respectively when considering that the latter numbers include animals with developmental defects that would not have hatched or survived in the wild and would therefore not have been included in the RAD-seq dataset.

## Discussion

In this study we report over 20 incidences of facultative parthenogenesis in marbled and Arizona striped whiptail lizards. By sequencing and assembling a highly contiguous *A. marmoratus* genome, we were able to refute automixis as the underlying mechanism in whiptail lizards. Instead, genome-wide homozygosity indicates a post-meiotic mechanism involving activation of embryonic development in unfertilized haploid oocytes. Even though FP whiptail lizards are largely comprised of homozygous diploid cells, a fraction of haploid cells persists through development and is readily detectable in young adults. Such mixoploidy and genome-wide homozygosity come at a price. In eight confirmed and 24 suspected cases of FP, development ceased prior to hatching and most FP animals that hatched showed congenital defects. Nevertheless, FP was observed at a rate of 1% and 5% in *A. arizonae* and *A. marmoratus*, respectively, and this occurred in the presence mating partners. Our findings indicate that FP is far more common in some vertebrate species than previously thought and the purifying selection associated with homozygosity may be an important force in generating additional resilience to counteract the effects of population bottlenecks and inbreeding depression.

The start of embryonic development is tightly coupled to fertilization in many vertebrates as the sperm entering the oocyte triggers a signaling cascade that is essential for the completion of female meiosis and initiation of cell division following karyogamy (51). This process has been mimicked in the laboratory by piercing frog oocytes with a needle to trigger the signal to complete meiosis and initiate replication and division in the haploid oocyte (52, 53). In zebrafish, homozygous embryos are routinely generated by fertilization of oocytes with UV-irradiated sperm, a treatment that destroys the paternal DNA (54). An oocyte treated in this manner will replicate the intact maternal genome in the absence of karyogamy. If the haploid oocyte is then subjected to heat shock treatment, cytokinesis is prevented resulting in a pseudodiploid oocyte that undergoes a second round of DNA replication followed by mitosis. Thus an entirely homozygous diploid embryo starts to develop (55).

Instead of two consecutive rounds of DNA replication without intervening mitosis at the start of embryonic development, our data indicate that one or multiple rounds of DNA replication and mitosis take place in the haploid oocyte prior to a skipped mitosis yielding a homozygous diploid cell followed by mixoploid development. Initiation of development in a haploid state may be conserved in avian species as unfertilized turkey eggs can yield embryos that contain 40% of haploid cells at blastoderm followed by a reduction to 1.3% within the blood of hatched birds (56). Depending on the tissue and ploidy distribution, the presence of haploid cells may contribute to abnormal development of specific tissues reported here and elsewhere (57, 58).

In addition to mixoploidy, genome-wide homozygosity constitutes another obstacle to normal development as each recessive deleterious allele is exposed in the hemizygous state. Indeed, arrested development and abnormal phenotypes are observed in FP whiptails, as well as in FP animals across many other species (16, 27, 59, 60). It is important to note though that some whiptails of FP origin developed normally, much like their sexually produced counterparts. At the population level, FP leads to a precipitous reduction in genetic diversity as only one set of alleles is inherited in the next generation. While FP could be an adaptive trait to bridge population bottlenecks when mate encounters are infrequent, small populations already rely heavily on inbreeding and FP further reduces the size of the gene pool (14, 25).

While FP can be considered the most extreme example of inbreeding, it is also the most extreme example of genetic purging as it eliminates most deleterious alleles in a single generation. FP in whiptail lizards and other species could therefore be considered a reproductive strategy, akin to mixed-mating systems in plants. While self-fertilization, or selfing, is reported in 10 to 15% of angiosperms, a low incidence of cross-fertilization, or outcrossing, cannot be ruled out (61). The ability to produce offspring via two different strategies provides a level of reproductive assurance in plants (62). Indeed, we see parallels to this in our own data in which female whiptails have produced offspring via both sexual reproduction and FP on separate occasions or simultaneously within a single clutch. Within plant species with mixed-mating, there are differences between the rates of selfing and outbreeding between populations and hypotheses as to why these differences occur include limited pollinator visitation and resource availability (63). To gain a better understanding of the origin and consequences of FP in whiptail lizards, it will be important to understand the triggers. It has been proposed that lack of or limited mate encounters triggers FP, but our data in combination with several other reports (10, 15, 25, 64) indicate that this is not the key trigger.

Our study adds whiptail lizards to a growing list of vertebrates capable of FP and establishes that it occurs alongside sexual reproduction in the presence of males. Using whole-genome sequencing, we unequivocally demonstrate that post-meiotic genome duplication is the underlying mechanism. One must now consider the possibility that FP is an adaptive trait and that low rates of successful FP contributes significantly to genome purification. Not only does FP reduce the frequency of deleterious alleles within a population, but it also provides reproductive assurance. Consequently, sexually mature FP offspring will have low genetic load and only pass on non-deleterious alleles to the next generation. Additional whole-genome sequencing data for species with documented FP will aid in the understanding the genetic basis, the propensity, and evolutionary significance of FP.

## Materials and Methods

### Animals

Animals used in this study were produced in the AAALAC International accredited Stowers Reptile and Aquatics Facility in compliance with protocols approved by the Institutional Animal Care and Use Committee. They descended from breeding stock collected in New Mexico under permit numbers 3199 and 3395 and Arizona under license number SP564133. Tissues used in the RAD-seq analyses were derived from samples collected under permits from the Arizona Game and Fish Department, California Department of Fish and Wildlife, New Mexico Department of Game and Fish, and the Secretariá de Medio Ambiente y Recursos Naturales, Dirección General de Fauna Silvestre of Mexico.

### Microsatellite analysis

DNA was extracted from tail samples for microsatellite genotyping as described in (65). PCR products were analyzed by capillary electrophoresis on a 3730 DNA Analyzer and data was analyzed using GeneMapper (v. 4.0). Primer information can be found in Table S7.

### Genome size estimation

The genome size of *A. marmoratus* was estimated by fluorescence-activated cell sorting (FACs), in which a standard curve correlating fluorescence intensity of DNA-bound propidium iodide with known genome sizes was generated using cells from fruit flies, zebrafish, and mouse, and then comparing fluorescent intensity with that of erythrocytes from *A. marmoratus*. Samples were stained using the Sigma PI staining preparation result in stained nuclei and ran on the Influx cytometer. PI fluorescence was collected using the PI Texas red detector with linear amplification and data analysis was performed in FlowJo and Microsoft Excel.

### DNA isolation, sequencing, and genome assembly for *A. marmoratus*

All genome sequencing libraries generated for the purpose of the *A. marmoratus* genome assembly were derived from the FP animal 8450. The liver tissue was first dissociated in a 10 mL Dounce homogenizer using the tight-fitting pestle and then processed using the Roche gDNA Isolation Kit (#11814770001, MilliporeSigma, St. Louis, MO, USA).

A short insert, high-coverage library was generated using the KAPA HTP kit (KK8234), with 1 µg of gDNA. The resulting library was size selected for fragments between 500-850 bp on a Pippin Prep (Sage Science). Two 40 Kb mate-pair libraries were generated by Lucigen from 1 µg of gDNA using the CviQl and BfaI restriction enzymes, respectively. Each library was sequenced on the Illumina MiSeq using the MiSeq Reagent Kit v2 (500 cycles). An additional three mate-pair libraries were generated, spanning distances of 5 Kb, 8 Kb, and 2-15 Kb, using the Illumina Nextera Mate-Pair Library Prep Kit and 1 µg of gDNA for each. Size selection used the Gel-Plus protocol with Pippin for the 5 and 8 Kb libraries, and the Gel-Free protocol for the 2-15 Kb library. All three libraries were pooled and sequenced on three separate RapidSeq flow cells on an Illumina HiSeq 2500. Chicago libraries were prepared from liver tissue sent to Dovetail Genomics (Dovetail Genomics LLC, Santa Cruz, CA, USA) to generate read pairs spanning distances up to 140 Kb and sequenced on an Illumina HiSeq 2500. The combined sequencing data was assembled at Dovetail Genomics using Meraculous and their in-house HiRise genome assembly algorithms to generate the *A. marmoratus* reference genome (AspMar1.0).

### Assessing assembly completeness

In order to assess the completeness of the *A. marmoratus* reference genome, we used BUSCO (v. 3.0.1) (66) with the vertebrate_od9 dataset containing 2,586 genes, with default parameters apart from changing the BLAST cutoff to 1e-6. We used BUSCO numbers generated in Gao *et al.* (67) for *Shinisaurus crocodilurus* and *Alligator mississippiensis* in Fig. S2*C*.

To perform a phylogenetic analysis, 1,333 shared “complete” single-copy orthologs were identified in the genomes of green anole (*Anolis carolinensis*, anoCar2), cow (*Bos taurus*, ARS-UCD1.2), dog (*Canis lupus familiaris*, CanFam3.1), zebrafish (*Danio rerio*, danRer10), chicken (*Gallus gallus*, galGal5), human (*Homo sapiens*, GRCh38.p13), mouse (*Mus musculus*, GRCm38.p6), medaka (*Oryzias latipes*, oryLat2), rat (*Rattus norvegicus*, Rnor_6.0), Argentine black and white tegu (*Salvator merianae*, HLtupMer3), western clawed frog (*Xenopus tropicalis*, Xenopus_tropicalis_v9.1), platyfish (*Xiphophorus maculatus*, X_maculatus-5.0-male). For each amino acid sequence a multiple sequence alignment was performed with MAFFT (v. 7.305) (68). The alignments were concatenated into a supermatrix of 1,112,277 amino acids. Phylogenetic tree topology was estimated using the Maximum Likelihood inference method using the pthreads version of RAxML (v. 8.2.11) and the PROTOGAMMAAUTO model for sequence evolution with 100 bootstrap replicates (69).

### Repeat identification

We quantified and annotated the repetitive DNA content within the *A. marmoratus* genome assembly by using the RepeatMasker pipeline on *A. marmoratus* scaffolds greater than or equal to 10 Kb in length. We first generated a de novo list of *A. marmoratus* repetitive elements using RepeatModeler (v. 1.0.11) (70). We then used these as input into RepeatMasker (v. 4.0.9) (71) using the NCBI/RMBLAST (v. 2.6.0+) search engine. Unclassified repeat element consensus sequences from the RepeatModeler output for each of the three lizards (*A. marmoratus, S. merianae,* and *A. carolinensis*) were compared to each other by identifying reciprocal best hits using BLAST (v. 2.9.0+).

### Whole-genome sequencing, reference genome alignment, and heterozygosity determination

Genomic DNA isolated from either liver or tail was prepared for sequencing using the KAPA HTP Library Preparation Kit (KK8234). Stock adapters were used from the Nextflex kit and barcodes were from BioScientific. All libraries were sequenced on the Illumina HiSeq 2500 platform. Whole-genome sequencing data was aligned to the *A. marmoratus* reference genome with BWA (v. 0.7.15) (72) and marked for duplicates with Picard (v. 1.119; https://broadinstitute.github.io/picard/). Because samples were sequenced over multiple lanes, the alignment files were merged subsequently, and another round of duplication marking was performed. The alignment files were realigned around small insertions and deletions with GATK (v. 3.5) (73). Data corresponding to lizard ID 122’s bam file was down-sampled to 33% of its original size using seqtk (v. 1.2-r94) to match the expected average genome coverage of the other samples.

The per position nucleotide profiles for each alignment were then generated using a combination of pysam (https://github.com/pysam-developers/pysam) and pysamstats (https://github.com/alimanfoo/pysamstats) to determine the heterozygosity at any genomic position.

### Transcriptome assembly and genome annotation

Two poly-A selected stranded RNA-sequencing libraries were generated with the TruSeq RNA Library Prep Kit v2 (RS-122-2001) and sequenced on an Illumina HiSeq 2500 for the purpose of an *A. marmoratus* transcriptome assembly. The first library was derived from a blood sample taken from a male animal, and the second library was derived from an embryo incubated at 28 °C and harvested 47-51 days post-egg deposition.

Trinity (v. 2.0.6) (74) was then used to generate an initial transcriptome assembly. The original reads were aligned to this transcriptome assembly using the Trinity companion script align_and_estimate_abundance.pl. Transcript isoforms with no read support were then filtered out and the remaining assembly was run through seqclean (https://sourceforge.net/projects/seqclean/). Evidence based annotations for the transcriptome assembly were generated using the MAKER2 pipeline (v. 2.31.8) (76). For MAKER2, the entire UniProtKB/Swiss-Prot database of proteins (Uniprot 2017) was used and the Repbase data base was used to mask repeats within the MAKER2 framework (77). Assigning putative function to the gene models was performed using BLAST (v. 2.6.0) and Interproscan (v. 5.13-52.0) (78).

Copy number estimation for the Vomeronasal 2 receptor 26 (Vmn2r26) genes was based on aligning the mouse ortholog (http://www.orthodb.org), to the *A. marmoratus* reference assembly using Exonerate (v. 2.4.0) (79) with a maximum intron size set to 20 Kb. Genes annotated as Vmn2r26 in the MAKER2 annotations were concatenated and aligned using Geneious (v. 10.1.3) (80) with default settings. The FastTree plugin (v. 1.0) was used to generate the phylogenetic tree from the alignment with default parameters.

### RAD-sequencing analysis

Double digest RAD-sequencing data was derived primarily from previous studies (35, 36, 81, 82). Fastq files were processed with Stacks (v. 2.62) using process_radtags and then samples from the same species were passed separately using denovo_map.pl. The average coverage for each sample was calculated and only samples that have at least a coverage of 20 were considered for subsequent analysis. For each passing sample, the heterozygosity was calculated for sites at average coverage.

### Giemsa staining of erythrocytes

Whole blood was collected aseptically from tails using acid citrate dextrose as anticoagulant. All samples were prepared immediately after collection. A 5 µL aliquot of the diluted blood sample was used to prepare a blood smear on a 25 x 75 x 1 mm microscope slide. Once dry, the slide was placed in 95% ethanol for 5 min. Giemsa stain (0.4%; Sigma, GS500) was applied liberally to cover the slide every 4 min for a total incubation time of 16 min. The prepared slides were imaged on an Axioplan2 imaging microscope equipped with a plan-apochromat 100x /1.40 Oil objective and an Axiocam HRc (color) camera (Zeiss). Micromanager (v. 1.4) software was used to acquire the images. The acquired images were then scored visually for the number of haploid, diploid, and binucleated erythrocytes.

### Feulgen staining of erythrocytes

Whole blood was collected aseptically from tails using acid citrate dextrose as anticoagulant. All samples were prepared immediately after collection. A 5 µL aliquot of the diluted blood sample was used to prepare a blood smear on a 25 x 75 x 1 mm microscope slide. Blood smears were treated with 10% neutral buffered formalin for 5 min at RT, then rinsed twice in distilled water. The slides were immersed in 5 M HCl for 30 min at RT, and then rinsed 2 times in distilled water. Slides were then immersed in Schiff’s reagent (Fisher Scientific #SS32-500) at RT for 15 to 30 min until nuclei were stained. The slides were transferred directly to bisulfite water that is prepared by dissolving 2.5 g of potassium metabisulfite in 500 mL of water and adjusting the pH to 4.0 with the addition of concentrated HCl. The bisulfite wash was repeated 3 times with 10-15 sec of agitation. The slide was then washed under running tap water for 2 min and dehydrated by incubating in 70% EtOH for 5 min and then 95% EtOH 5 min. The preparations were cleared in xylene before mounting.

### Imaging of ovaries and germinal vesicle

Germinal vesicles and acquisition of image stacks by confocal microscopy were performed as described in (33). Ovary images were acquired with a Leica M205FA dissection microscope with a planar 0.63X objective. Micromanager (v. 1.4) software was used to acquire images.

## Data availability

All raw sequencing data pertaining to the AspMar1.0 genome assembly are available at the National Center for Biotechnology Information under project accession number PRJNA360150. All other whole-genome and RNA sequencing data can be found under PRJNA980964. RAD-seq data are under: PRJNA827355, PRJNA707030, PRJNA762930, and PRJNA1016487. Code and raw microsatellite data used for analysis are available at https://github.com/baumannlab/Aspi_FP.

## Acknowledgments

We are grateful for the husbandry staff at the Stowers Institute for Medical Research (SIMR) especially Rick Kupronis and the team of dedicated reptile technicians, David Jewell, Alex Muensch, Jillian Schieszer, Christina Piraquive, and Kristy Winter, for outstanding husbandry and herpetocultural skills. We also thank the members of the New Mexico Department of Game and Fish and the Arizona Game and Fish Department for their support. We thank Veronica Cloud for help with RNA extractions. We thank the molecular biology, flow cytometry, and advanced microscopy core facilities at SIMR for their support and the members of the Emergent AI Center at Johannes Gutenberg University Mainz (JGU) for excellent IT support and helpful discussions. We acknowledge computing time granted on the supercomputer MOGON II at JGU as part of NHR South-West (nhrsw.de), as well as the Institute of Molecular Biology Bioinformatics Core for computing resources. This work was funded in part by the Howard Hughes Medical Institute, SIMR, and an Alexander von Humboldt Professorship awarded to P.B. at JGU. This work also received support from the National Science Foundation grant DEB-1754350 and the GenEvo RTG funded by the Deutsche Forschungsgemeinschaft (German Research Foundation) – GRK2526/1 – Projectnr. 40.7023052 and the Institute for Quantitative and Computational Biosciences (IQCB) in Mainz. We thank Charles (Jay) Cole, Uri García-Vázquez, Rick Kupronis, Daniel Lara-Tufiño, Norma Manríquez-Moran, Maximiliano Monroy, Priscilla Neaves, Adrián Nieto Montes de Oca, Harry Taylor, Robert Thomson, and Carol Townsend for assistance with fieldwork and specimen collection. We also thank the Natural History Museum of Los Angeles County, the UTEP Biodiversity Collections, Jay Cole and the American Museum of Natural History, the Museum of Vertebrate Zoology, Randy Klabacka and the Auburn University Museum of Natural History, and the LSU Museum of Natural Science for providing tissue loans in support of this research.

---

**Figure S1.**
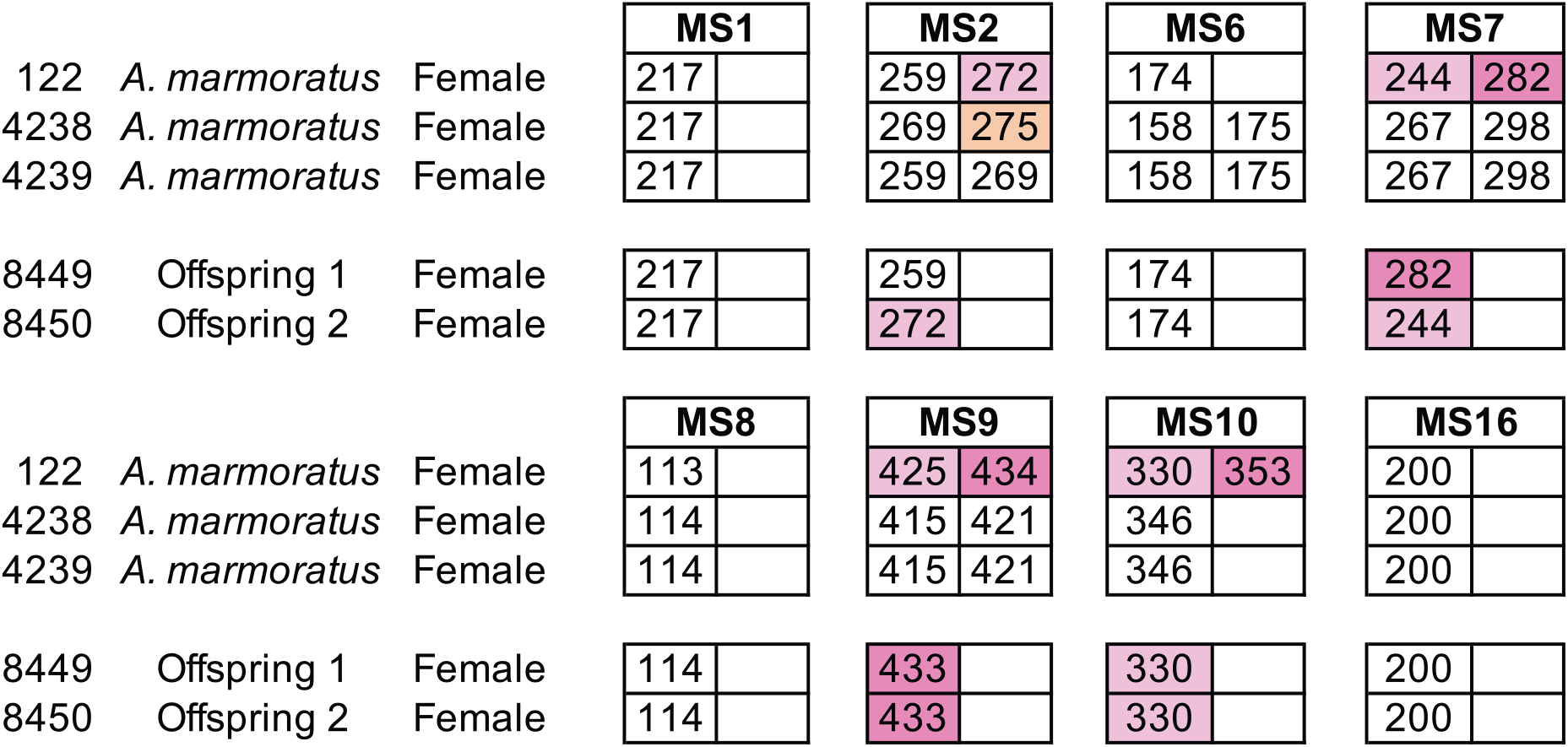
Microsatellite analysis of eight loci for the three female *Aspidoscelis marmoratus* within the enclosure and the two offspring hatched in January 2009. Only alleles that are unique to a potential mother are colored. Single nucleotide differences in size may indicate allele differences but may be binning artifacts and therefore are not scored as different alleles. The two offspring were homozygous for all eight markers. At three markers (MS7, MS9, MS10), both offspring inherited alleles only found in animal 122.

**Figure S2.**
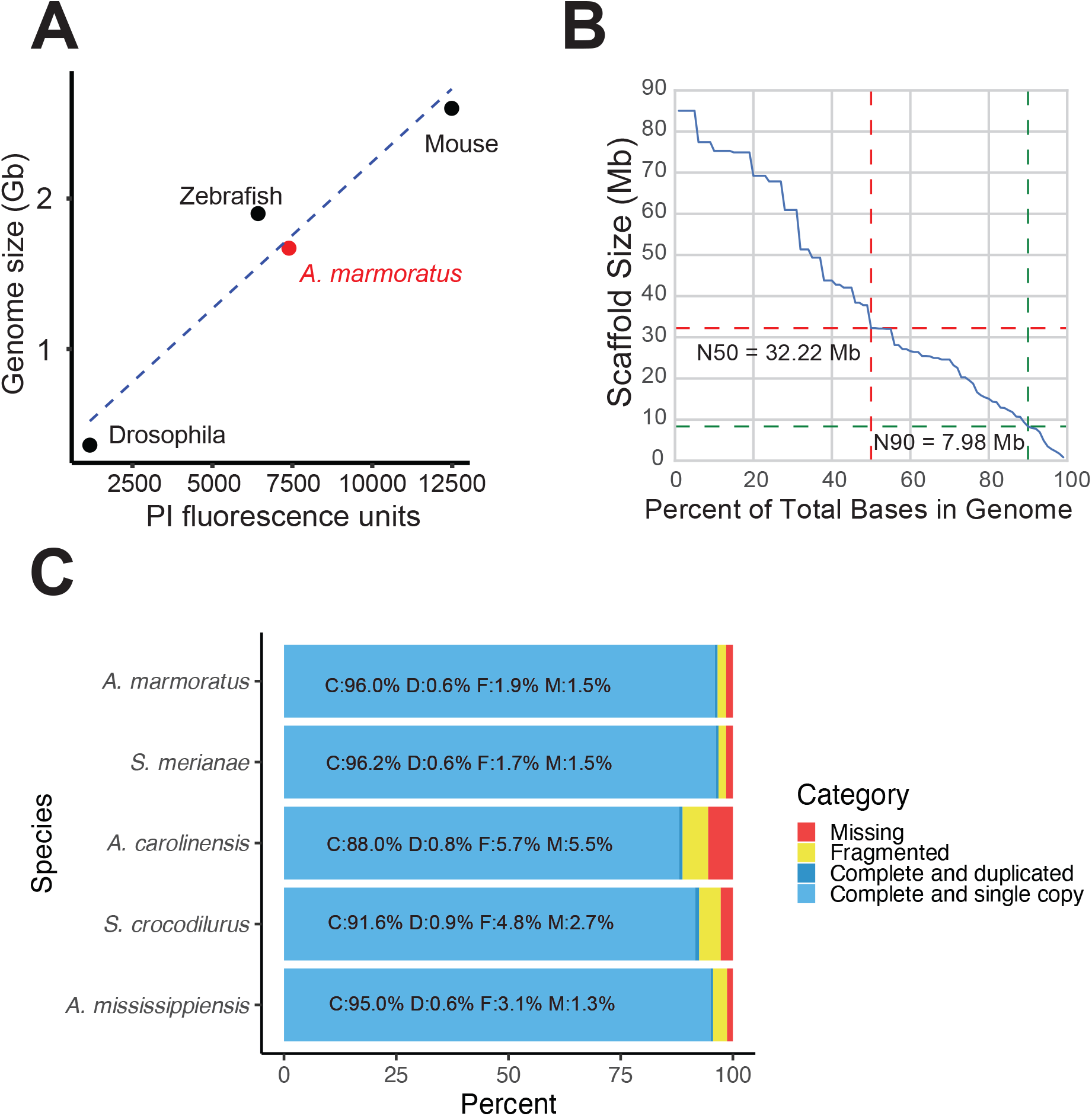
Genome assembly of *Aspidoscelis marmoratus*. **(A)** Genome size estimation using fluorescence-activated cell sorting (FACs). The standard curve was generated using known genome sizes from fruit flies, zebrafish, and mouse, and then comparing fluorescent intensity with that of erythrocytes from *A. marmoratus*. **(B)** N(x)% plot shows the ordered scaffold lengths plotted against the cumulative fraction of the genome. Dashed lines mark the N50 and N90 values. **(C)** BUSCO analysis of 2,586 conserved vertebrate coding genes for *A. marmoratus*, *Salvator merianae*, *Anolis carolinensis*, *Shinisaurus crocodilurus*, and *Alligator mississippiensis*. Values for *S. crocodilurus* and *A. mississippiensis* were taken from *Gao et al.* (1).

**Figure S3.**
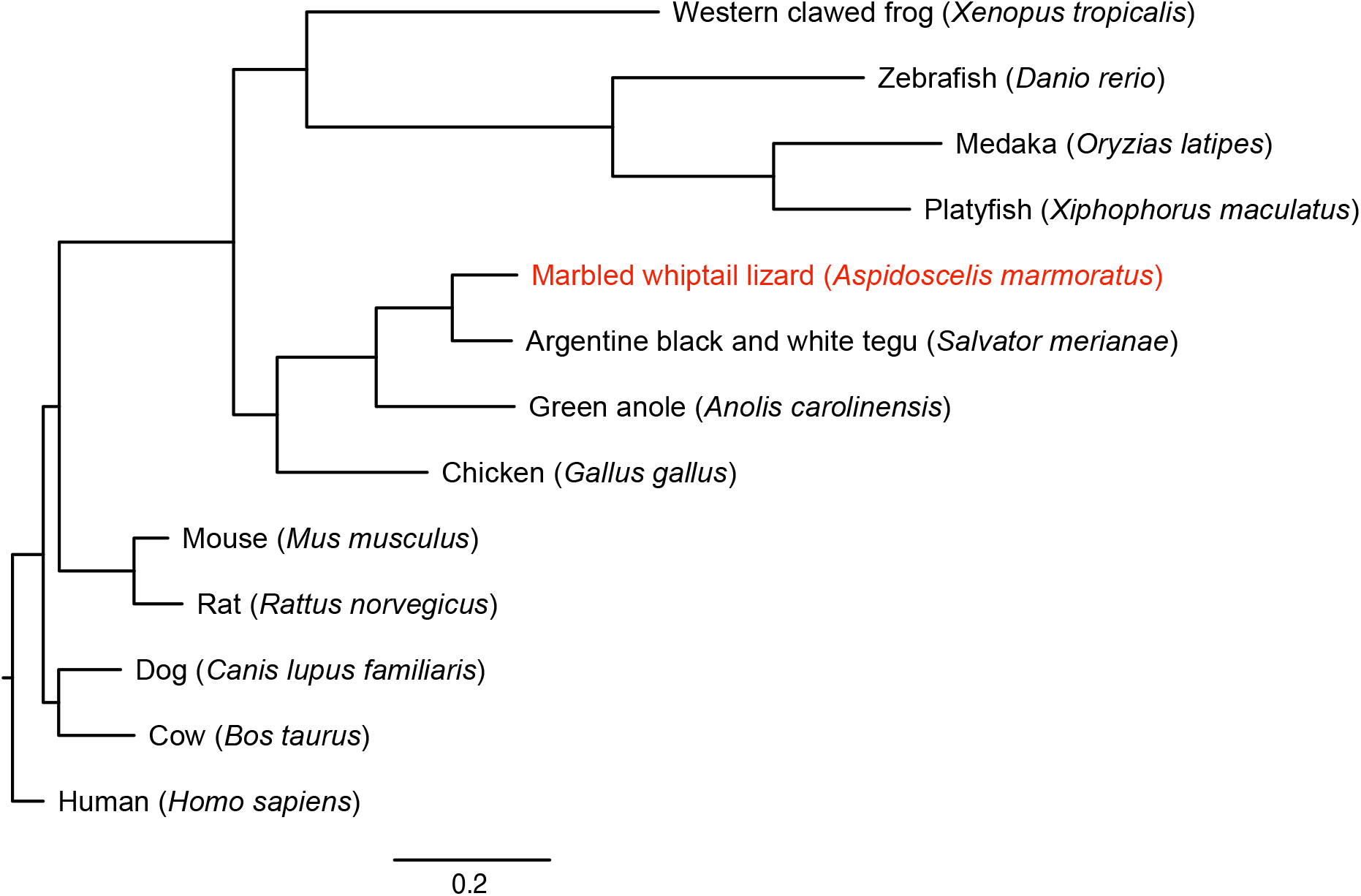
Maximum likelihood tree for 13 vertebrate genomes, based on 1,333 single-copy BUSCOs detected across all species analyze.

**Figure S4.**
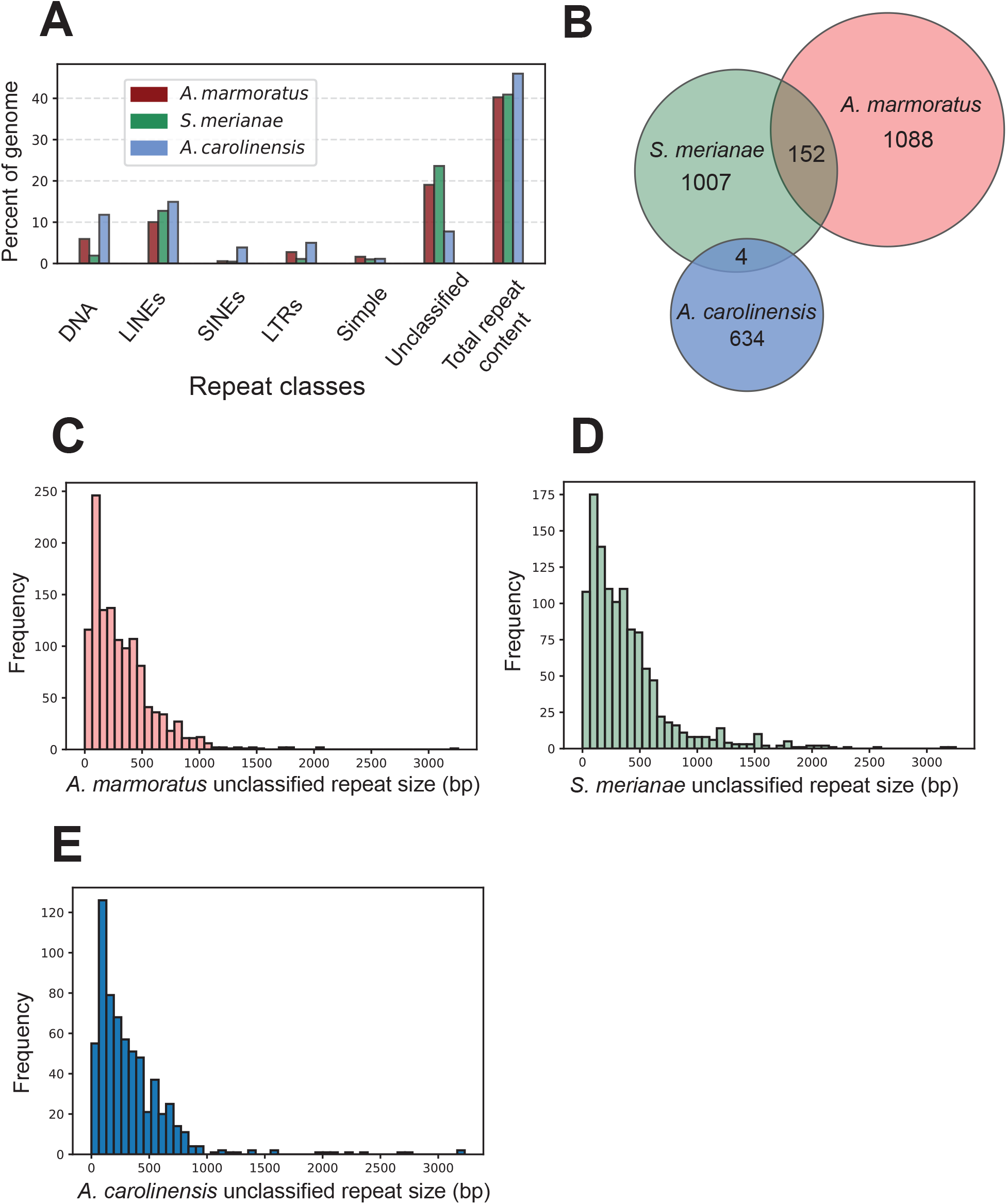
**(A)** Repetitive DNA content in *Aspidoscelis marmoratus*, *Salvator merianae*, and *Anolis carolinensis* genomes separated by repeat classes as defined by RepeatMasker. **(B)** Overlap of unclassified repetitive elements between the three genomes as in (A). **(C-E)** Distribution of the sizes of unclassified repetitive elements, respectively, for *A. marmoratus*, *S. merianae*, and *A. carolinensis*.

**Figure S5.**
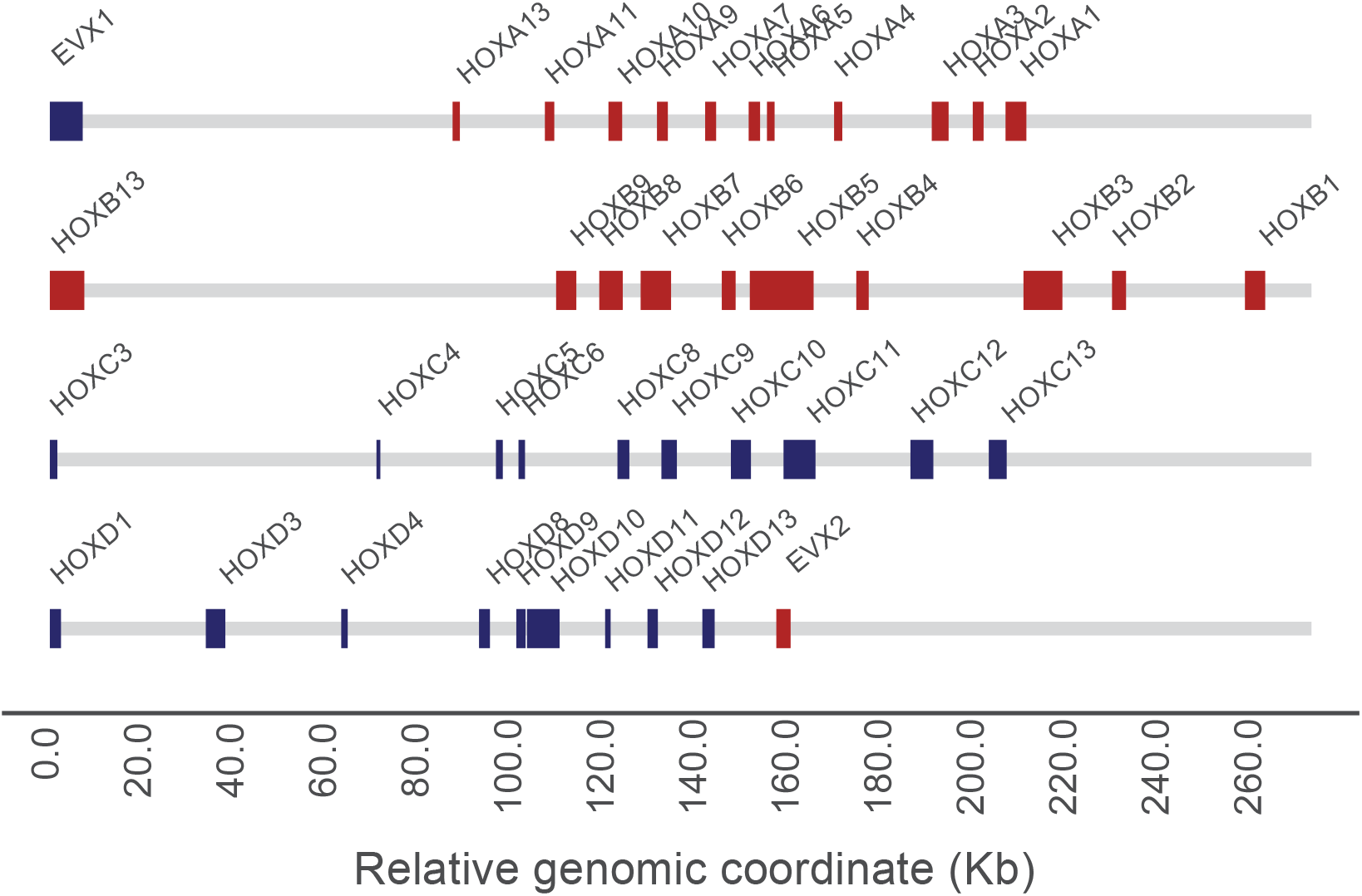
*Aspidoscelis marmoratus* HOX gene clusters. Red blocks indicate a region of homology was found on the sense strand, blue indicates the antisense strand. EVX1 and EVX2 are also shown, due to their relevance to HOX cluster evolution.

**Figure S6.**
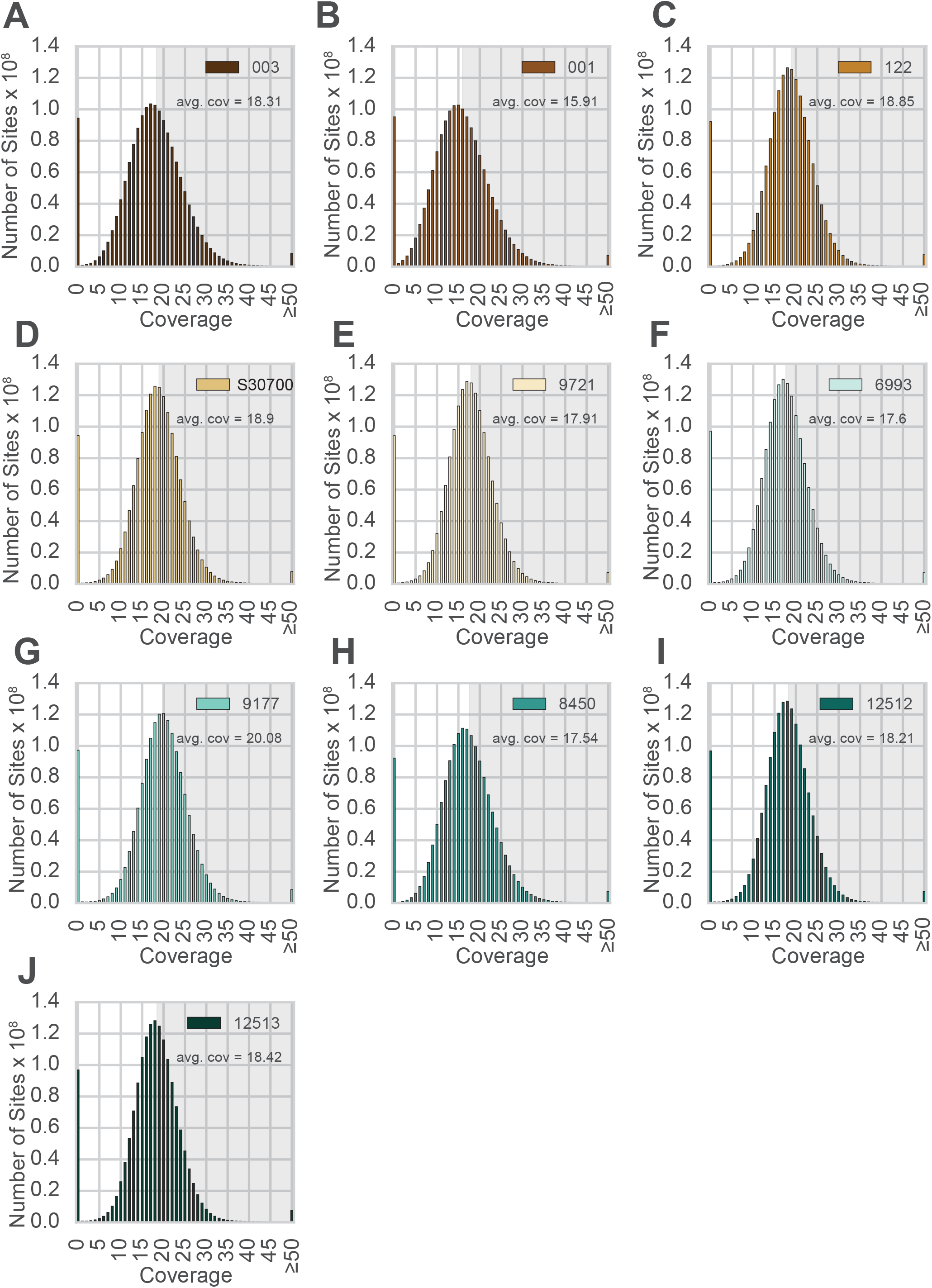
**(A-J)** Distribution of coverage for all animals sequenced. Animal IDs are shown in the top right corner in each panel. The distributions were generated by examining the coverage at every position in the reference assembly. Regions with coverage greater than the mean are shaded in grey in each panel.

**Figure S7.**
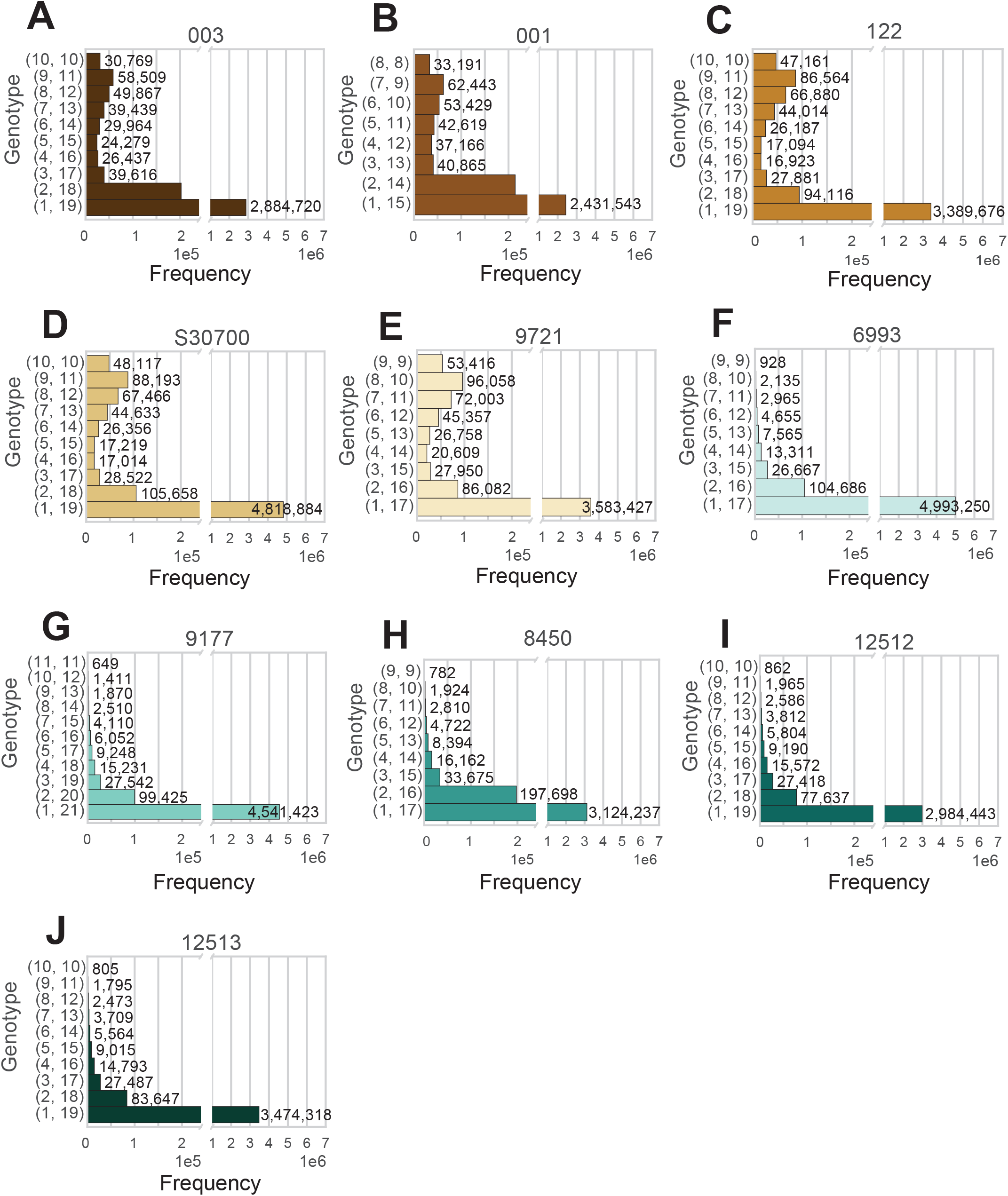
**(A-J)** Analysis of all positions in the genome at average coverage, for which reads support exactly two alleles. The two numbers to the left of each row indicate how many reads support Frequency each allele. Animal IDs are shown above each panel.

**Figure S8.**
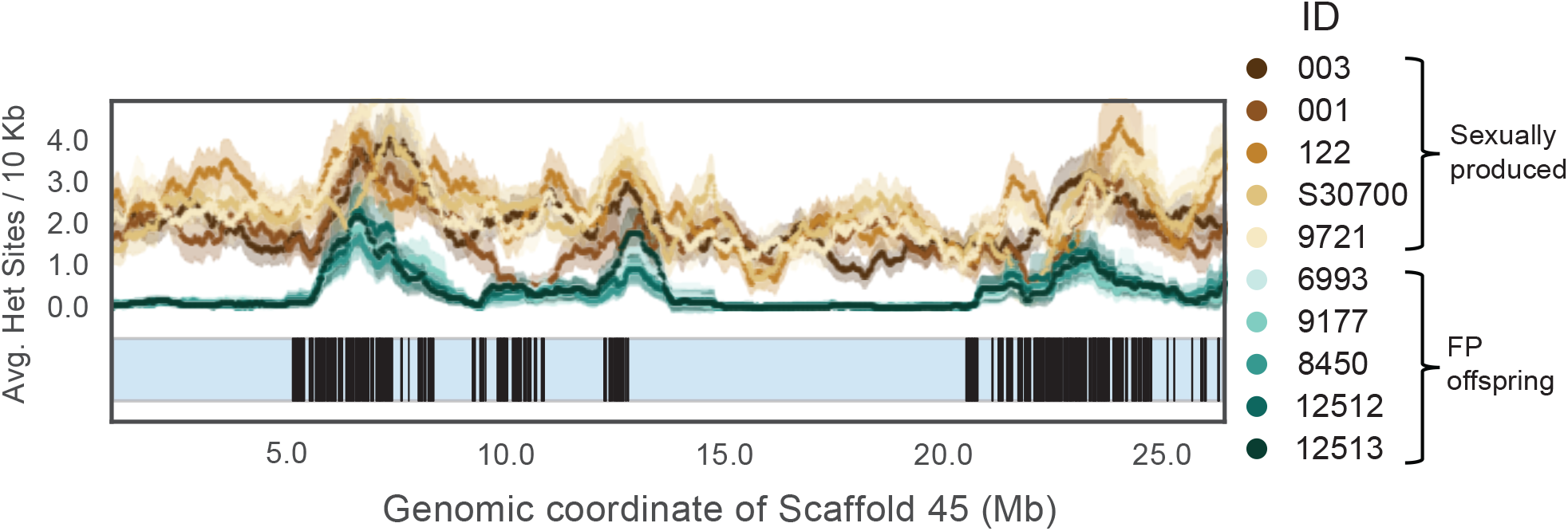
A 1 Mb sliding window for Scaffold 45. A scaffold ideogram is shown in blue and coordinates of 167 annotated V2r26 homologs are shown in black.

**Figure S9.**
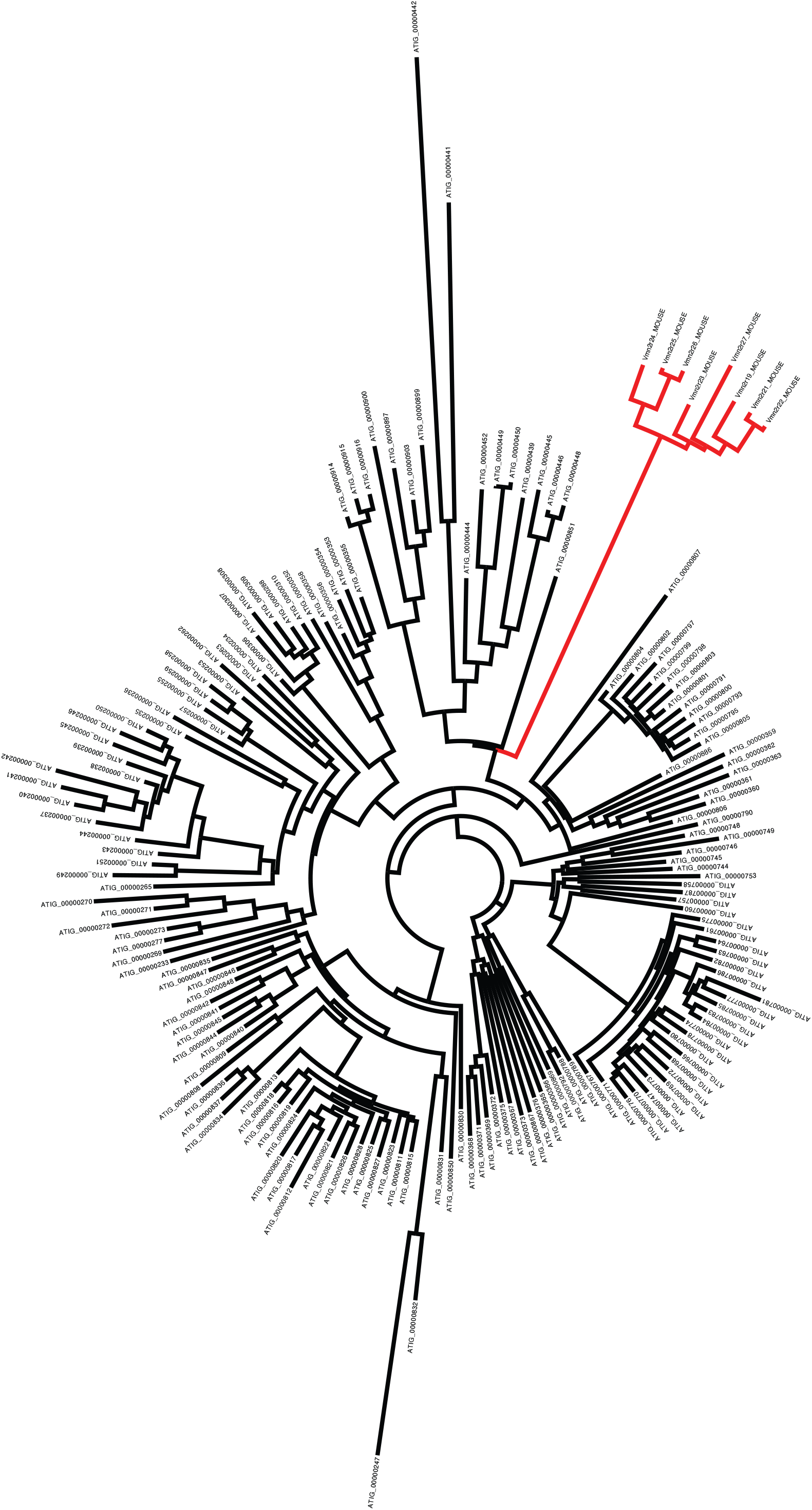
Phylogenetic tree showing evolutionary distance between mouse V2Rs (red branches) and *Aspidoscelis marmoratus* homologs (black branches). There are 10 additional copies of Vmn2r26 homologs found on Scaffold 45 not identified by MAKER2 (177 total), while genome-wide, we annotated 323 copies. Phylogenetic analysis of the 323 copies with 8 other mouse vomeronasal 2 receptors show that the mouse receptors form their own distinct clade from *A. marmoratus* and there is substantial sequence diversity among the *A. marmoratus* homologs. A manual search for additional copies of Vmn2r26 resulted in 478 further hits. Vmn2r26 belongs to a family of receptors known as V2Rs and this estimate on the number of Vmn2r26 homologs ranks *A. marmoratus* as one of the species with the largest expansion of V2Rs (2, 3).

**Figure S10.**
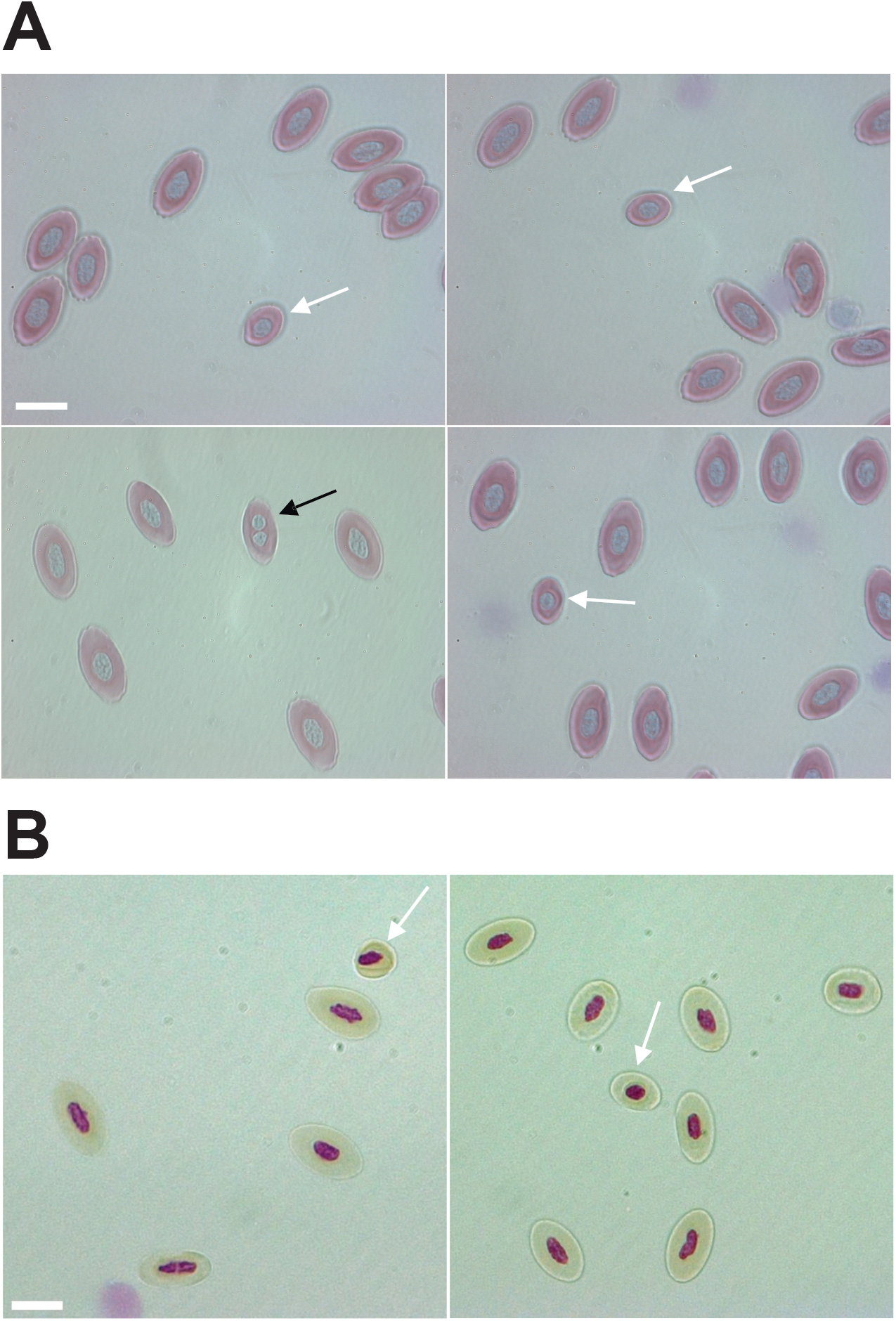
**(A)** Giemsa staining of erythrocytes from a one-year-old *Aspidoscelis marmoratus* produced by facultative parthenogenesis. Diploid (n = 1661), smaller haploid (white arrow, n = 17), and binucleated (black arrow, n = 1) cells are evident. Scale bar corresponds to 10 µm. **(B)** Feulgen staining of erythrocytes from a one-year-old *A. arizonae* produced by facultative parthenogenesis. Diploid (n = 130) and smaller haploid (white arrow, n = 2) cells are evident. Scale bar corresponds to 10 µm.

**Figure S11.**
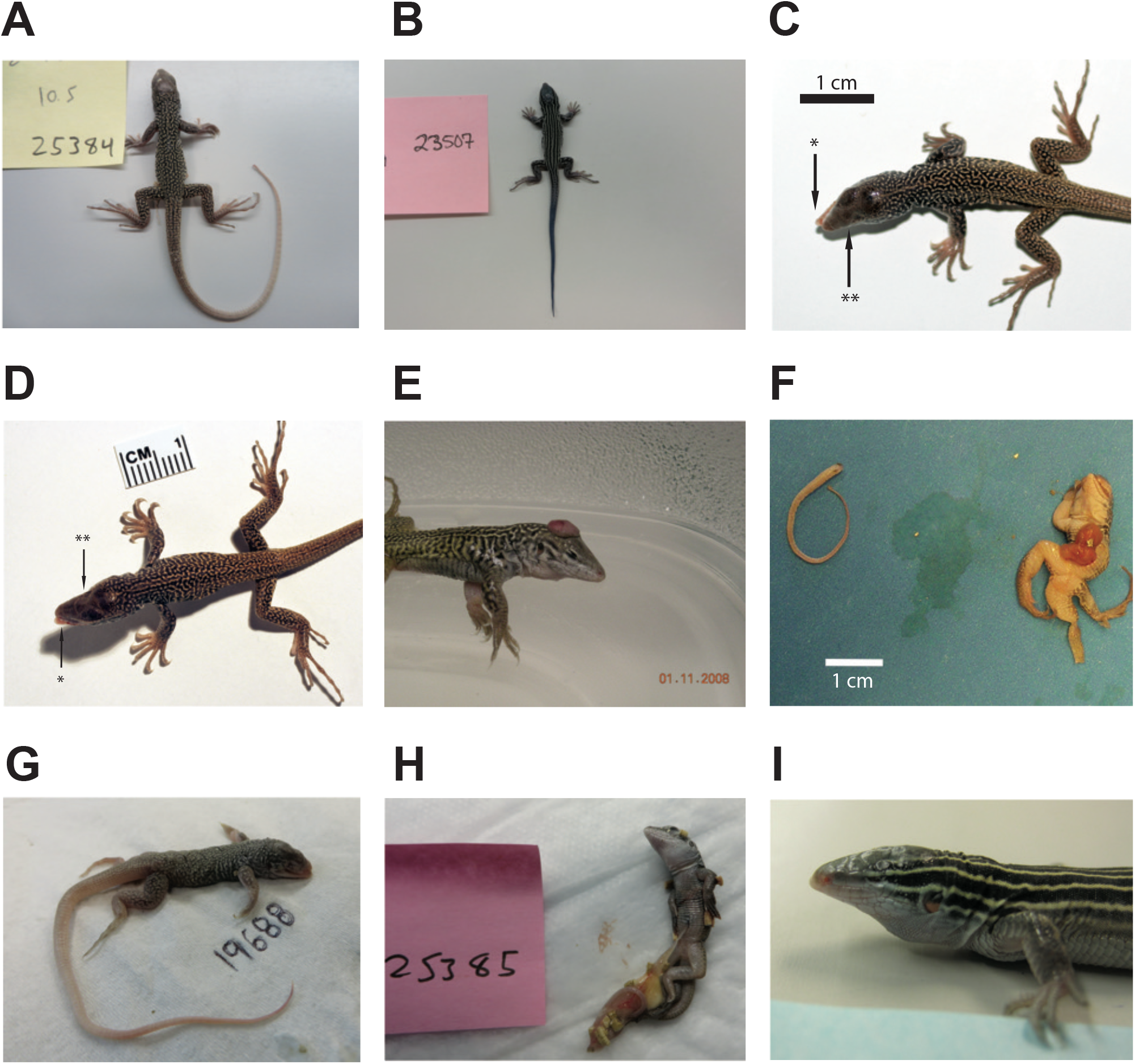
Animals produced by facultative parthenogenesis. **(A)** *Aspidoscelis marmoratus* 25384. No abnormal phenotypes noted. **(B)** *Aspidoscelis arizonae* 23507. No abnormal phenotypes noted. **(C)** *A. marmoratus* 12512. Misalignment of jaws (*). Agenesis of left eye (**). **(D)** *A. marmoratus* 12513. Misalignment of jaws (*). Agenesis of right eye (**). **(E)** *A. marmoratus* 6993. Animal was cut from egg with exposed brain. **(F)** *A. marmoratus* 8394. Did not hatch. Missing a leg, failure of abdomen closure leading to exposed organs, face abnormalities, and hunched back. **(G)** *A. marmoratus* 19688. Did not hatch. Multiple craniofacial deformities. **(H)** *A. marmoratus* 25385. Partially emerged from egg with egg yolk still attached. **(I)** *A. arizonae* 16216. Agenesis of left eye.

**Figure S12.**
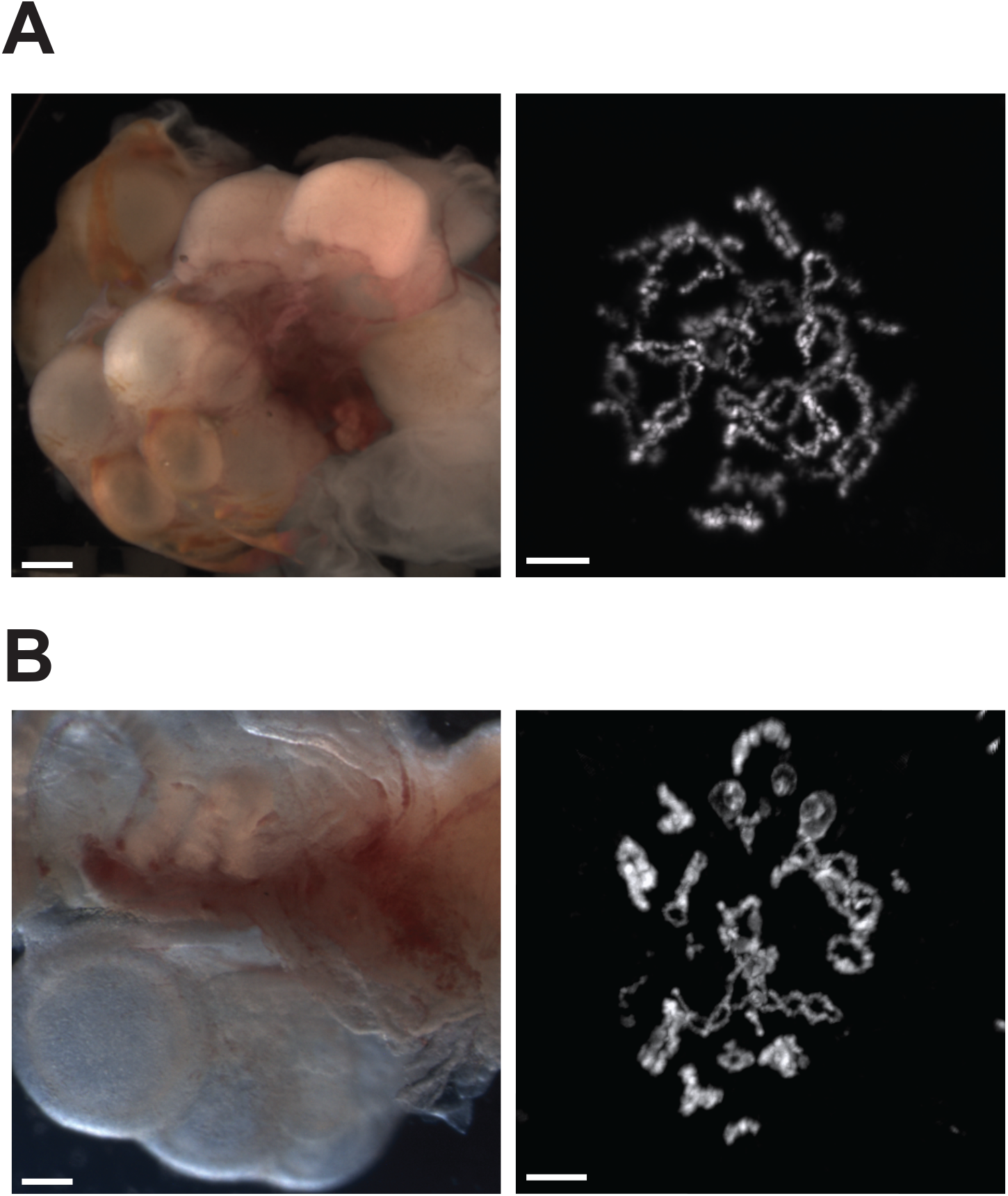
No differences between ovaries and germinal vesicles of *Aspidoscelis marmoratus* produced by facultative parthenogenesis or fertilization. **(A)** Ovary (left) and diplotene stage germinal vesicle (right) from *A. marmoratus* produced by facultative parthenogenesis. The lizard ovary image (ID 8450) is taken with a Leica M205FA dissection microscope. Scale bar corresponds to 1 mm. The germinal vesicle (ID 8449) is stained with DAPI. Scale bar corresponds to 10 µm. **(B)** Ovary (left) and diplotene stage germinal vesicle (right) from a sexually produced *A. marmoratus*. The lizard ovary image (ID 13103) is taken with a Leica as described previously. Scale bar corresponds to 1 mm. The germinal vesicle (ID 5359) is stained with Acridine Orange (0.01%). Scale bar corresponds to 10 µm.

**Figure S13.**
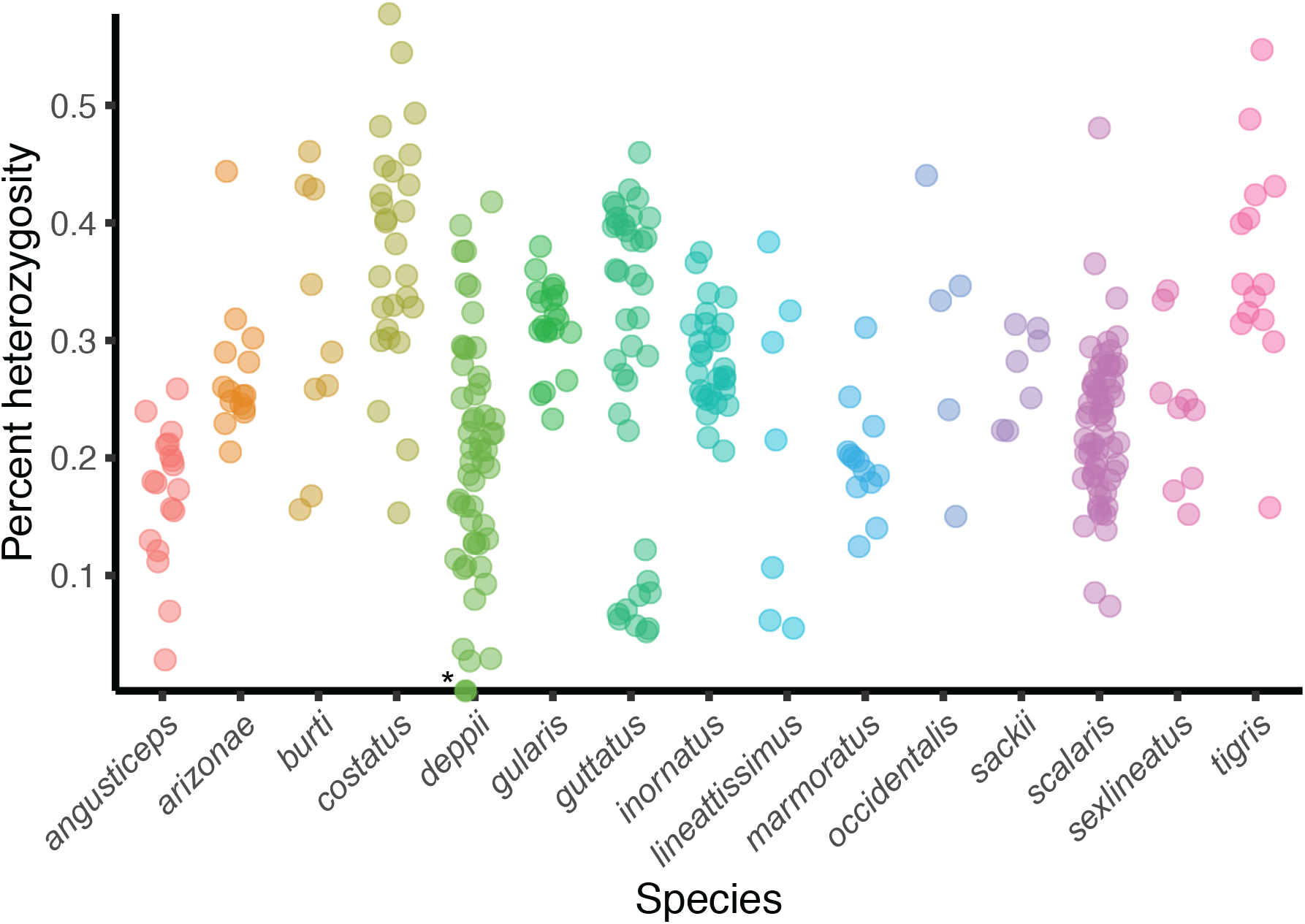
Facultative parthenogenesis in whiptail lizards collected in nature. Percent heterozygosity estimates from RAD-seq for 321 whiptail lizards from 15 species. All individuals had an average coverage of at least 20. Each point is an individual, and percent heterozygosity was calculated only for sites where the coverage is equal to the average sequencing coverage. The heterozygosity of one *Aspidoscelis deppii* individual (ID LDOR30, marked with *) is far less than that observed for individuals of the same species (Rosner’s Test for Outliers, p < 0.001), having only one called heterozygous position. Species (sample size): *angusticeps* (18), *arizonae* (15), *burti* (9), *costatus* (27), *deppii* (52), *gularis* (20), *guttatus* (38), *inornatus* (28), *lineattissimus* (7), *marmoratus* (13), *occidentalis* (5), *sackii* (7), *scalaris* (60), *sexlineatus* (9), *tigris* (14).

**Table S1.** *A. marmoratus* genome assembly statistics.

| Statistic | Meraculous | HiRise | Units |
| --- | --- | --- | --- |
| N50 | 1600688 | 32220929 | bp |
| N90 | 413554 | 7979424 | bp |
| max_scaffold | 11507399 | 85027298 | bp |
| mean_scaffold | 188287.18 | 428523.466 | bp |
| median_scaffold | 1703 | 1450 | bp |
| min_scaffold | 1000 | 927 | bp |
| number of scaffolds | 8705 | 3826 | bp |
| perc_A | 27.06 | 27.04 | % |
| perc_C | 20.14 | 20.12 | % |
| perc_G | 20.13 | 20.13 | % |
| perc_N | 5.64 | 5.67 | % |
| perc_T | 27.04 | 27.04 | % |
| perc_genome_covered_by_scaffolds_greater_than_100kb | 97.99 | 99.46 | % |
| perc_genome_covered_by_scaffolds_greater_than_10kb | 99.28 | 99.63 | % |
| perc_genome_covered_by_scaffolds_greater_than_1mb | 69.83 | 98.47 | % |
| scaffolds_greater_than_100Mb | 0 | 0 |  |
| scaffolds_greater_than_100kb | 1588 | 133 |  |
| scaffolds_greater_than_10Mb | 2 | 45 |  |
| scaffolds_greater_than_10kb | 2076 | 223 |  |
| scaffolds_greater_than_1Mb | 545 | 90 |  |
| total bases | 1639039918 | 1639530780 | bp |
| non_n_GC% | 42.67 | 42.67 | % |
| non_n_AT% | 57.33 | 57.33 | % |

**Table S2.**
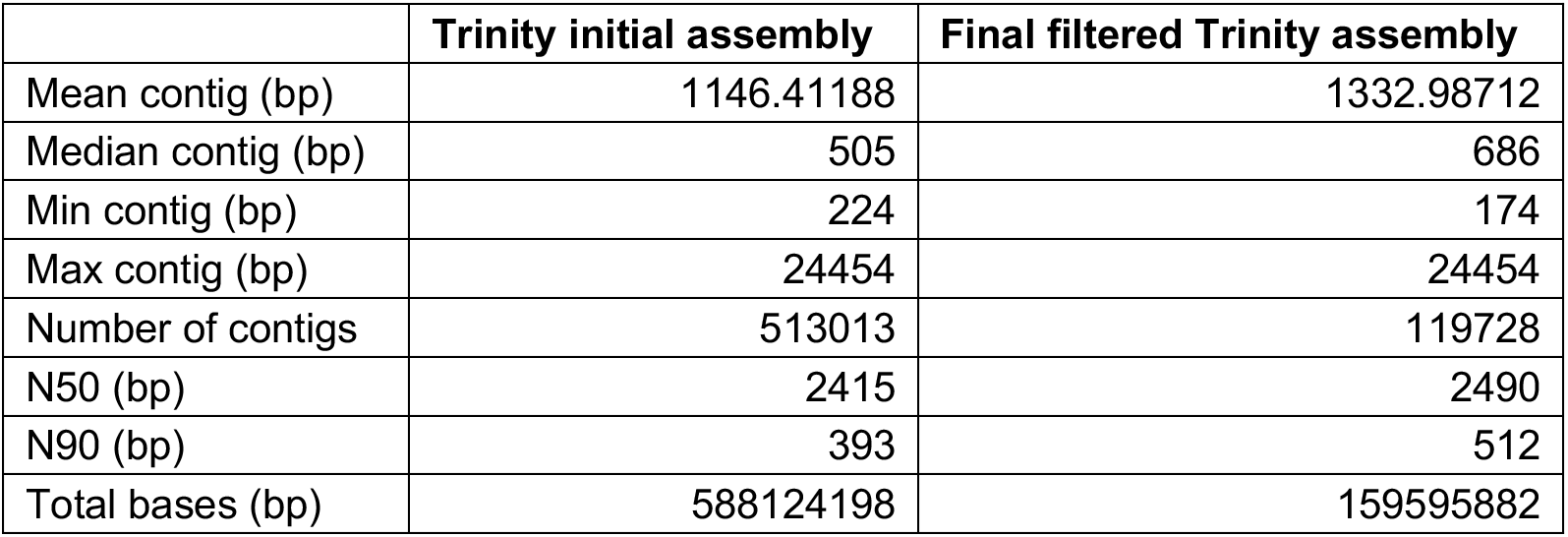
Trinity assembly statistics.

**Table S3.** MAKER2 summary.

| Feature | Source | Count |
| --- | --- | --- |
| CDS | MAKER2 | 222450 |
| exon | MAKER2 | 164669 |
| five_prime_UTR | MAKER2 | 22402 |
| gene | MAKER2 | 25856 |
| intron | MAKER2 | 189732 |
| mRNA | MAKER2 | 44461 |
| three_prime_UTR | MAKER2 | 21729 |

**Table S4.**
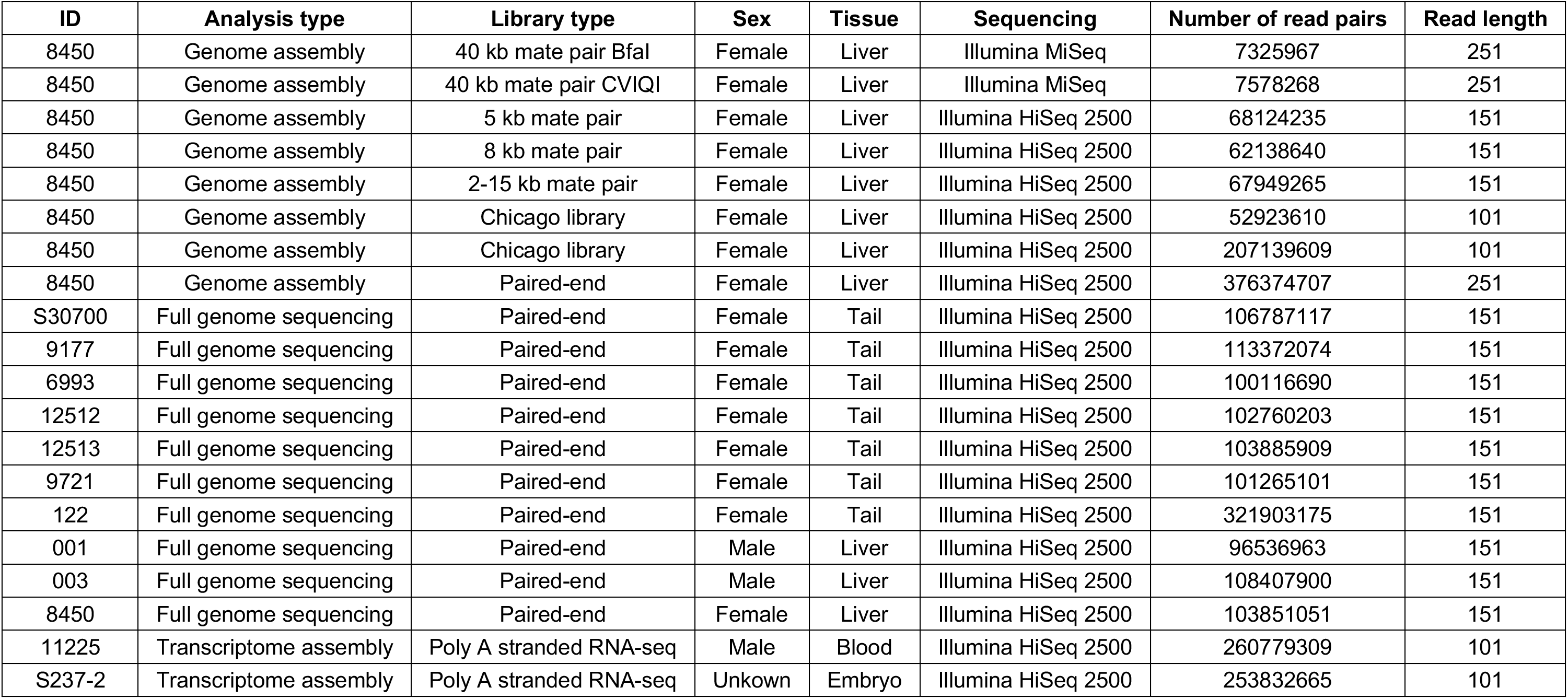
Information for all *A. marmoratus* animals sequenced.

**Table S5.** Whole genome sequencing alignment statistics.

| **Animal ID** | **12512** | **12513** | **S30700** | **6993** | **9177** | **9721** | **001** | **003** | **122** | **8450** |
| --- | --- | --- | --- | --- | --- | --- | --- | --- | --- | --- |
| **total** | 2.1E+08 | 2.12E+08 | 2.18E+08 | 2.04E+08 | 2.32E+08 | 2.07E+08 | 1.97E+08 | 2.21E+08 | 6.56E+08 | 2.12E+08 |
| **secondary** | 4371764 | 4293355 | 4427220 | 4120929 | 4761643 | 4246124 | 3932037 | 4507406 | 12408444 | 4081669 |
| **supplementary** | 0 | 0 | 0 | 0 | 0 | 0 | 0 | 0 | 0 | 0 |
| **duplicates** | 1829281 | 1841233 | 1848550 | 2314935 | 2169035 | 2151550 | 1523198 | 1283723 | 14760637 | 1368644 |
| **mapped** | 2.07E+08 | 2.09E+08 | 2.15E+08 | 2.01E+08 | 2.29E+08 | 2.04E+08 | 1.93E+08 | 2.18E+08 | 6.5E+08 | 2.08E+08 |
| **paired in sequencing** | 2.06E+08 | 2.08E+08 | 2.14E+08 | 2E+08 | 2.27E+08 | 2.03E+08 | 1.93E+08 | 2.17E+08 | 6.44E+08 | 2.08E+08 |
| **read1** | 1.03E+08 | 1.04E+08 | 1.07E+08 | 1E+08 | 1.13E+08 | 1.01E+08 | 96536963 | 1.08E+08 | 3.22E+08 | 1.04E+08 |
| **read2** | 1.03E+08 | 1.04E+08 | 1.07E+08 | 1E+08 | 1.13E+08 | 1.01E+08 | 96536963 | 1.08E+08 | 3.22E+08 | 1.04E+08 |
| **properly paired** | 1.88E+08 | 1.9E+08 | 1.95E+08 | 1.82E+08 | 2.07E+08 | 1.85E+08 | 1.84E+08 | 2.06E+08 | 5.97E+08 | 1.96E+08 |
| **with itself and mate mapped** | 2.02E+08 | 2.04E+08 | 2.09E+08 | 1.95E+08 | 2.22E+08 | 1.99E+08 | 1.88E+08 | 2.13E+08 | 6.36E+08 | 2.02E+08 |
| **singletons** | 1212491 | 1504914 | 1467678 | 1679077 | 1523933 | 1408129 | 1072182 | 1099052 | 1912936 | 993280 |
| **with mate mapped to a different chr** | 12512434 | 12380462 | 12772150 | 11827242 | 13772076 | 12316058 | 3751076 | 5431708 | 34786180 | 5413508 |
| **with mate mapped to a different chr (mapQ>=5)** | 7450173 | 7412852 | 7583394 | 7060453 | 8164436 | 7358325 | 2149235 | 3193570 | 21253145 | 3196901 |

**Table S6.** Animals confirmed by microsatellite analysis to be of FP origin.

| Species | Animal ID | Clutch ID | Laid-Begin | Laid-End | Hatch | Notes |
| --- | --- | --- | --- | --- | --- | --- |
| *A. marmoratus* | 6993 | X48 | 21-Oct-07 | 22-Oct-07 |  | Animal was cut from egg, alive, with brain exposed from skull |
| *A. marmoratus* | 8377 | J68 | 21-Oct-08 | 25-Oct-08 |  |  |
| *A. marmoratus* | 8394 | P68 | 29-Oct-08 | 2-Nov-08 |  | Missing a leg, organs exposed at abdomen, face abnormalities, and hunched back |
| *A. marmoratus* | 8449 | M69 | 18-Nov-08 | 22-Nov-08 | 25-Jan-09 | Appeared to have balance issues; hatched with substance still attached to abdomen |
| *A. marmoratus* | 8450 | M69 | 18-Nov-08 | 22-Nov-08 | 25-Jan-09 | Appeared to have balance issues |
| *A. arizonae* | 8677 | K72 | 7-Feb-09 | 10-Feb-09 |  |  |
| *A. marmoratus* | 9070 | V78 | 12-Jun-09 | 16-Jun-09 |  |  |
| *A. marmoratus* | 9177 | F80 | 30-Jul-09 | 3-Aug-09 | 10-Oct-09 |  |
| *A. marmoratus* | 12512 | L131 | 26-Apr-12 | 30-Apr-12 | 8-Jul-12 | Deformed jaw; missing left eye |
| *A. marmoratus* | 12513 | L131 | 26-Apr-12 | 30-Apr-12 | 8-Jul-12 | Deformed jaw, missing right eye |
| *A. arizonae* | 16215 | B187 | 7-Mar-14 | 11-Mar-14 | 2-May-14 | Right side of skull seems to have stunted growth compared to left side, lower mandible protrudes out slightly, right eye seems slightly smaller than left |
| *A. arizonae* | 16216 | B187 | 7-Mar-14 | 11-Mar-14 | 2-May-14 | Missing left eye, malformed jaw, shortened torso |
| *A. marmoratus* | 19687 | T237 | 19-May-15 | 23-May-15 | 1-Aug-15 |  |
| *A. marmoratus* | 19688 | T237 | 19-May-15 | 23-May-15 |  | Multiple craniofacial deformities |
| *A. arizonae* | 23304 | Q288 | 11-Sep-16 | 15-Sep-16 | 15-Nov-16 |  |
| *A. arizonae* | 23507 | D291 | 11-Oct-16 | 15-Oct-16 | 15-Dec-16 |  |
| *A. marmoratus* | 24514 | L303 | 12-Jan-17 | 16-Jan-17 |  |  |
| *A. arizonae* | 25339 | I313 | 3-Mar-17 | 7-Mar-17 |  | Deformed snout, missing left eye |
| *A. marmoratus* | 25384 | R313 | 5-Mar-17 | 9-Mar-17 | 6-Jun-17 |  |
| *A. marmoratus* | 25385 | R313 | 5-Mar-17 | 9-Mar-17 | 6-Jun-17 | Found partially emerged from egg, barely moving, had large mass of egg yolk still attached |
| *A. arizonae* | 29606 | E364 | 27-Apr-18 | 1-May-18 | 3-Jul-18 |  |
| *A. arizonae* | 29607 | E364 | 27-Apr-18 | 1-May-18 | 3-Jul-18 | Congenital shortened torso; hatched alive and died immediately |
| *A. marmoratus* | 30050 | A369 | 29-Jun-18 | 3-Jul-18 |  | No eyes |

**Table S7.**
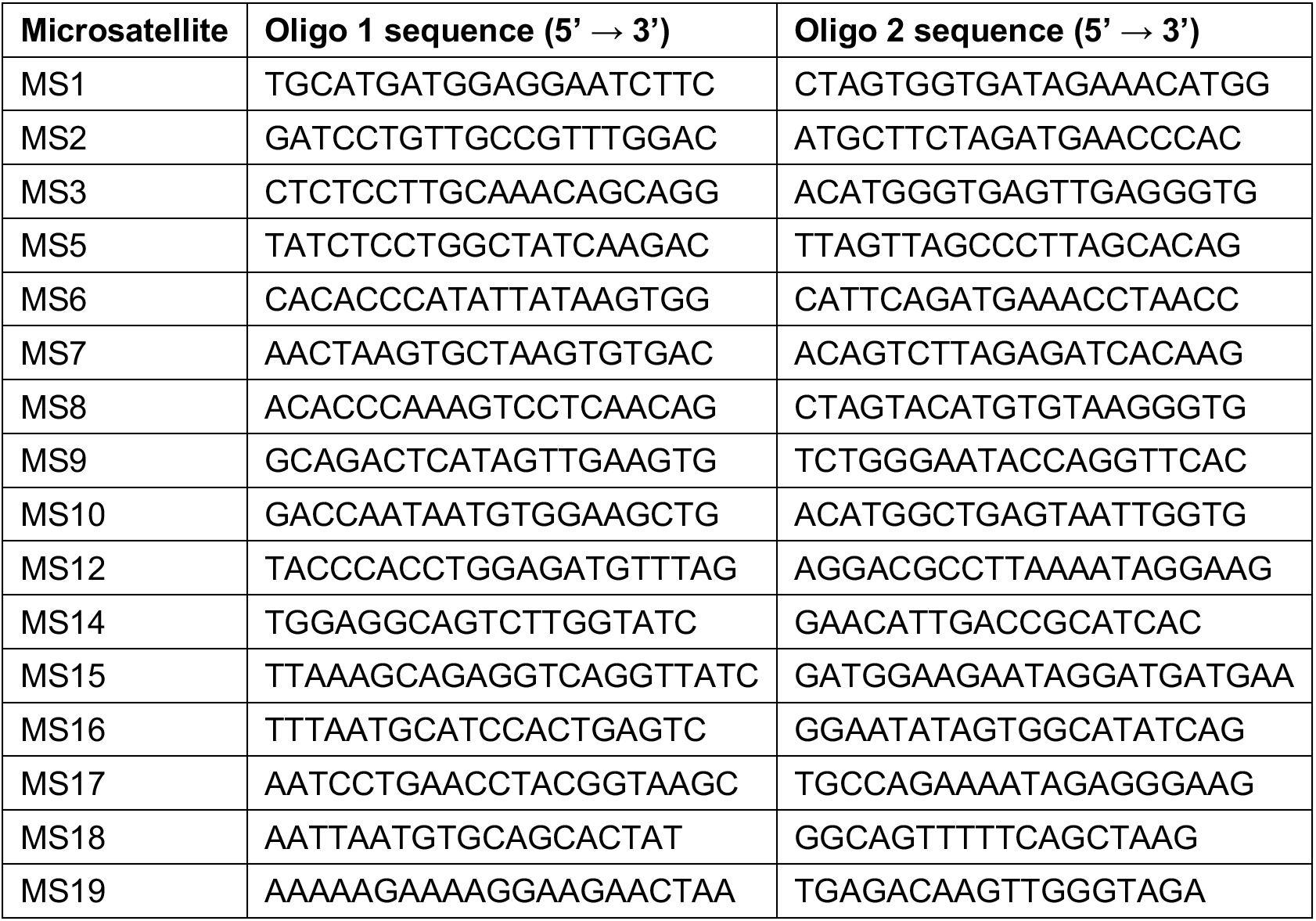
Microsatellite primer information.

## Footnotes

* The taxonomy of the genus *Aspidoscelis* has undergone frequent revisions and the maternal ancestor of the obligate parthenogenetic species *A. neomexicanus* was formerly known as the subspecies *A. tigris marmoratus* and the male ancestor as the subspecies *A. inornatus arizonae* (35). For the purpose of this manuscript we follow the taxonomic conclusions by Barley *et al.* (36).

